# The gut microbiota metabolite Urolithin A mitigates JAK signaling to suppress cytokine–mediated autoimmune diseases

**DOI:** 10.64898/2026.05.08.723914

**Authors:** Shijin Geng, Rong-Chun Tang, Hengxiang Yu, Ao Zhang, Shuang-Shuang Yu, Yunxuan Zhou, Lan Zhang, Jun Zhang

## Abstract

Aberrant activation of type I interferon (IFN-I) is closely related to the development of autoimmune diseases. The metabolic regulation of cytokine signaling is essential for immune homeostasis. In this study, we characterized Urolithin A(UA), a natural gut-derived metabolite, as an inhibitor of Janus kinase (JAK) signaling. UA was found to broadly dampen JAK phosphorylation and the downstream signaling induced by cytokines such as type I interferons (IFN-I), type II interferons (IFN-II), and interleukin-6 (IL-6). UA can directly bind to JAK1 JH1 domain and treatment with UA attenuated autoimmune pathogenesis in *Trex1*-KO mice, IMQ-induced SLE and psoriasis models. Our findings unveil that UA is an anti-inflammatory metabolite that promotes immune homeostasis and could be used to treat inflammatory and autoimmune diseases.

## Introduction

The human gut microbiota produces a wide range of bioactive metabolites that act as important links between microbial ecology and host immunity ^1,2^. Over the past decade, certain small-molecule metabolites among these microbial byproducts, have emerged as potent immunomodulators capable of fine-tuning inflammatory responses without inducing broad immunosuppression. Accumulating evidence has revealed that these microbial metabolites exert anti-inflammatory effects through diverse mechanisms, including 1. direct modulation of pattern recognition receptor (PRR) and immune signaling ^3^, 2. epigenetic regulation of immune cell gene expression ^4,5^, 3. metabolic reprogramming of inflammatory pathways ^6^, and 4. direct engagement of host receptors, including AhR, GPCRs and nuclear receptors ^7–9^, *etc*.

The Janus kinase-signal transducer and activator of transcription (JAK–STAT) signaling pathway acts as a fundamental conduit in immune regulation, transducing signals from numerous cytokines, interferons, and growth factors to orchestrate gene expression programs that govern cell survival, differentiation, and effector function ^10,11^. Dysregulation of this pathway, whether due to gain-of-function mutations (GOFs), aberrant cytokine signaling, or epigenetic modifications, has been strongly implicated in the pathogenesis of autoimmune diseases, including rheumatoid arthritis (RA), systemic lupus erythematosus (SLE), psoriasis and inflammatory bowel disease (IBD) ^12^. For instance, the JAK1 V658F mutation in the pseudokinase domain (JH2), analogous to the classic JAK2 V617F, disrupts the autoinhibitory function of the JH2 domain, which may mimic a state of persistent cytokine stimulation ^13^.

Similarly, the STAT1 GOF mutations (for example, R274W and T385M) heightened prolonged STAT1 phosphorylation in response to interferons, which not only predisposes individuals to chronic mucocutaneous candidiasis due to impaired Th17 immunity, but also creates a pervasive IFN-signature that dramatically increases susceptibility to a spectrum of organ-specific autoimmune pathologies ^14,15^. While JAK inhibitors (e.g., tofacitinib, baricitinib) have demonstrated clinical success in treating autoimmune disorders, their broad impact on multiple cytokine pathways is associated with an increased risk of infections and other adverse events ^16,17^. Furthermore, resistance mechanisms and the incomplete efficacy in certain patient groups highlight the need for alternative strategies to modulate JAK–STAT signaling more precisely ^17^. Emerging evidence suggests that natural compounds, such as microbial metabolites, may fine-tune this pathway with fewer side effects.

Urolithin A (UA) is a gut microbiota-derived terminal metabolite of ellagitannins and ellagic acid (EA) generated by specific bacterial taxa such as *Enterocloster asparagiformis*, *Enterocloster bolteae*, *Enterocloster citroniae* and *Enterocloster pacaense* ^18^. UA has attracted considerable interest as a clinically advanced postbiotic because it enhances mitochondrial function, activates mitophagy and autophagy, and has shown beneficial effects in preclinical models and human studies of metabolic and age-related conditions ^19–22^. In immune cells, UA has been reported to suppress NF-κB activation and inflammasome assembly, suggesting broader anti-inflammatory potential ^23,24^. However, whether UA directly interfaces with canonical cytokine signaling pathways, particularly the JAK–STAT axis that underpins many autoimmune diseases, has not been addressed. Considering the growing global burden of autoimmune disorders and the limitations of current immunosuppressive therapies (e.g., high toxicity, partial efficacy, and relapse risks), exploring immunoregulatory properties of UA could provide novel, safer effective treatment options.

In this study, we characterized the immunoregulatory role of UA in cytokine mediated signaling and elucidated its therapeutic potential in autoimmune disease models. This is the first report to demonstrate that UA directly targets JAK1, highlighting its therapeutic value in autoimmune disorders. It also provides evidence how a dietary metabolite modulates inflammatory response, implicating the interplay between microbial metabolism and immune homeostasis. This work may inspire the dietary microbiome-based therapies, which may offer a natural, low-toxicity alternative to conventional immunosuppressants.

## Results

### UA is a potent inhibitor of IFN-I induced signaling pathway

While the anti-inflammatory properties of UA have been documented ^19,23,24^, its potential role in cytokine-mediated signaling pathways remains largely unexplored. Given the broad physiological and pathological significance of IFN-I signaling, particularly regarding its dysregulation in autoimmune diseases, we sought to investigate whether UA modulates IFN-I-triggered responses, with the aim of evaluating its therapeutic potential in interferonopathies and related autoimmune diseases.

To determine whether UA modulates IFN-I signaling, we first isolated primary peritoneal macrophages from C57BL/6 mice after thioglycollate induction (Figure 1A). CCK-8 assays showed that UA had no measurable cytotoxicity at concentrations up to 20 µM, which were the range adopted in subsequent experiments (Figure 1B). Pretreatment with UA significantly attenuated the induction of classical interferon-stimulated genes (ISGs), including *Ifit1* and *Ifit2*, in a time-dependent manner following stimulation with murine IFN-α (Figure 1C). At the protein level, immunoblotting revealed that UA markedly suppressed IFN-α–induced phosphorylation of JAK1, TYK2, STAT1, STAT2, and STAT3 without altering the total levels of these signaling components (Figure 1, D and E). To assess the activity of UA in a physiological setting, the mice were administered UA orally once daily for seven days, followed by intravenous challenge with murine IFN-α (Figure 1F). Quantitative PCR analysis of tissue samples collected 4 hours post stimulation demonstrated that UA pretreatment significantly reduced transcripts of ISGs including *Isg15*, *Ifit1*, *Ifit2*, and *Oas1* in the liver, kidney, and heart compared with DMSO-treated controls (Figure 1G).

**Figure 1.**
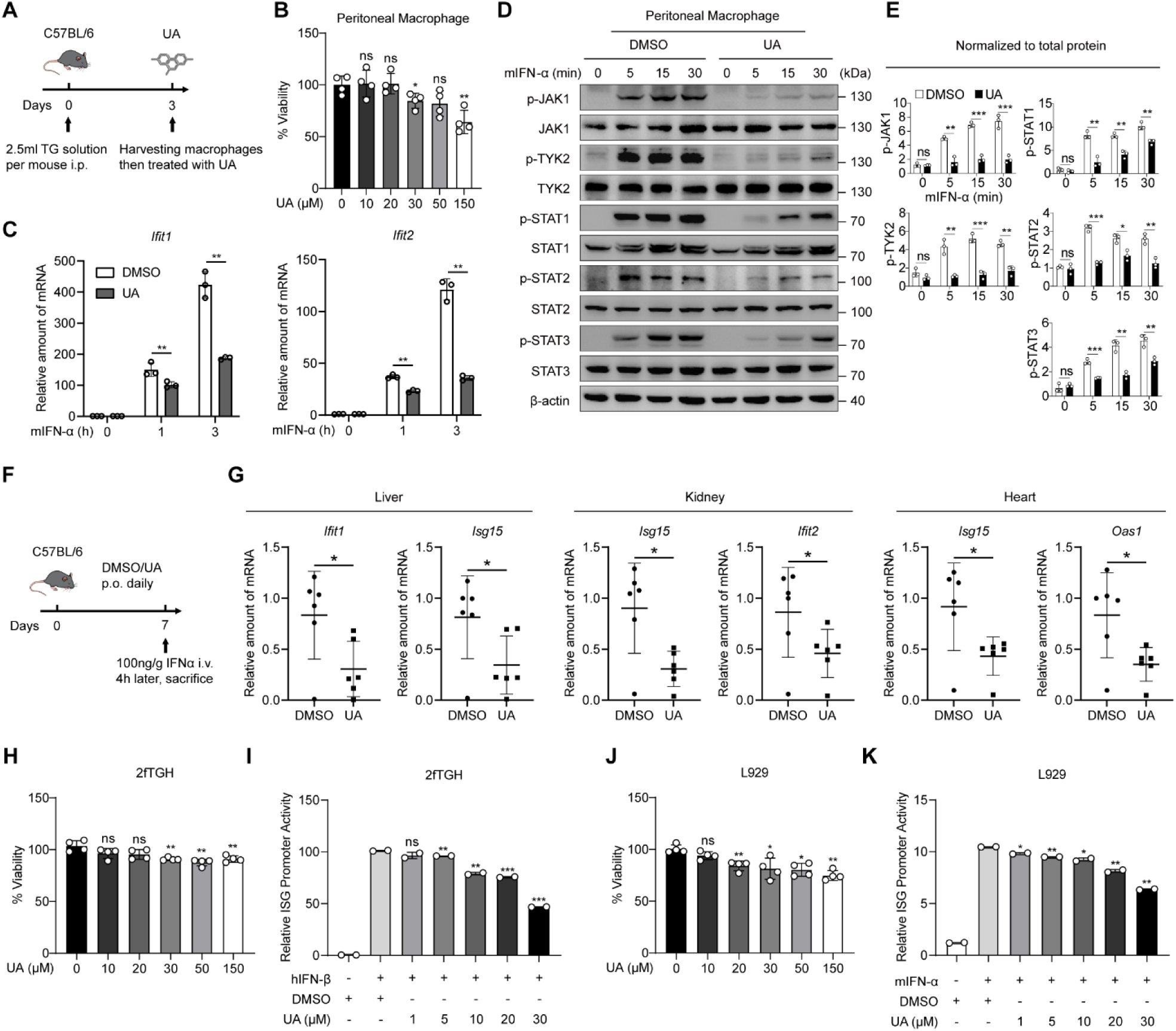
UA inhibits IFN-I signaling in primary peritoneal macrophages and in ISRE reporter cell lines. **A**, Schematic of experimental workflow for isolation of thioglycollate elicited primary peritoneal macrophages and subsequent *in vitro* UA treatment. Mice received 2.5 ml 3% (w/v) thioglycollate solution intraperitoneally, macrophages were harvested at day 3, which were then collected for qPCR and immunoblot analysis. **B**, Cell viability of primary peritoneal macrophages after 24 h exposure to UA at the concentrations indicated in the panel, measured by CCK-8 assay. Data are shown relative to DMSO control. **C**, qPCR analysis of *Ifit1* and *Ifit2* mRNA levels in primary peritoneal macrophages pretreated with 20µM UA for 24 h and stimulated with mouse IFN-α (10ng/mL) for the indicated times. **D**, Immunoblot analysis of primary peritoneal macrophages after UA pretreatment and IFN-α stimulation. Cells were incubated with 20µM UA or DMSO for 24 h, then stimulated with 10ng/mL mouse IFN-α for the indicated times. **E**, Densitometric quantification of phosphorylated proteins normalized to the corresponding total proteins in **D**, plotted for the indicated times. **F**, Schematic of the *in vivo* treatment regimen. C57BL/6 mice received oral UA(100mg/kg) or DMSO once daily for seven days, followed by intravenous injection of mouse IFN-α (100ng/g). Tissues were collected four hours after IFN-α administration. **G**, qPCR analysis of interferon-stimulated genes (ISGs) in liver, kidney and heart from mice treated with UA or DMSO before mouse IFN-α challenge. Each symbol represents one mouse (n = 6 per group). **H**,**J**, Cell viability of human 2fTGH cells and murine L929 cells, respectively, after 24 h exposure to UA at the concentrations indicated, measured by CCK-8 assay. **I**,**K**, ISRE luciferase reporter assays in 2fTGH and L929 cells, respectively. Cells were pretreated with UA at the concentrations indicated, then stimulated with human IFN-β (2fTGH) or mouse IFN-α (L929). Luciferase activity is presented relative to the stimulated DMSO control. Statistical significance is indicated: ns = no significance, \**P* < 0.05, \*\**P* < 0.01, \*\*\**P* < 0.001.

We further evaluated its effects using interferon-stimulated response element (ISRE) luciferase reporters in human 2fTGH and murine L929 cells. Consistent with the primary macrophage data, UA exhibited no cytotoxicity in either cell line within the tested dose range (Figure 1, H and J). Luciferase assays demonstrated dose-dependent inhibition of IFN-I–induced ISRE activation, using human IFN-β in 2fTGH cells and mouse IFN-α in L929 cells (Figure 1, I and K).

Together, these results consistently establish UA as a potent inhibitor of IFN-I signaling, supporting its further exploration as a candidate immunomodulatory agent.

### UA suppresses JAK–STAT activation to attenuate IFN-I signaling independent of autophagy

Based on the observed inhibition of IFN-I signaling by UA in primary macrophages and *in vivo* models, we next sought to determine the underlying molecular mechanism of this suppression. To delineate the precise step at which UA acts within the IFN-I signaling cascade, we systematically examined its effect on JAK–STAT pathway activation in HEK293T, HeLa and mouse embryonic fibroblast (MEF) cells. As was observed in primary mouse macrophages, where UA inhibited IFN-α–induced JAK–STAT phosphorylation (Figure 1D), this inhibitory effect was conserved in all cell lines tested. The cells were pretreated with 10 µM UA (a concentration that showed no cytotoxicity by CCK-8 assay; Supplemental Figure 1, A–C) prior to stimulation with species-matched IFN-I. Immunoblot analysis revealed that UA pretreatment significantly suppressed the phosphorylation of both JAK1 and TYK2, as well as their downstream STAT1, STAT2, and STAT3 in these tested cell types throughout the 60-min time course, without affecting total protein levels (Figure 2, A, D and G). Quantification by densitometric scanning of immunoblot bands, normalized to the corresponding total protein, is shown in Figure 2, C, F and I and corroborates the inhibitory effect of UA on JAK–STAT phosphorylation. The consistent inhibition observed at the phosphorylation level of JAK1 and TYK2—the most upstream kinases in the pathway—indicates that UA targets the initial activation step of IFN-I signaling. In accordance with the diminished JAK–STAT signaling, qPCR confirmed the reduced expression of canonical ISGs (*ISG15*, *IFIT1* and *OAS1* in human cells, and *Isg15*, *Ifit1* and *Oas1* in MEFs) after UA treatment (Figure 2, B, E and H).

**Figure 2.**
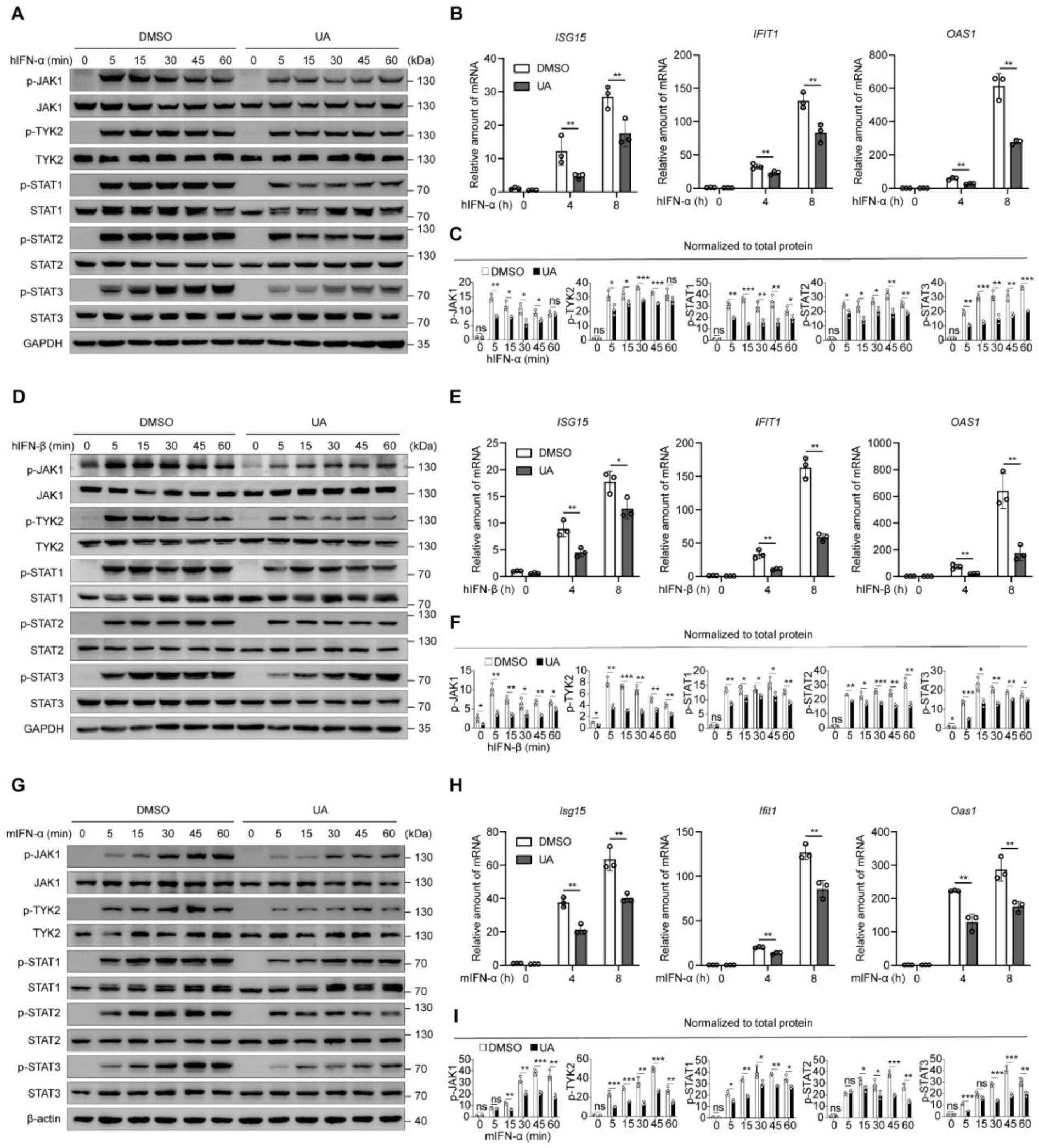
UA suppresses IFN-I signaling across human and murine cell lines. **A**,**D**,**G**, Immunoblot analysis of HEK293T, HeLa and MEF cells pretreated with 10 µM UA for 24 h and then stimulated with 10ng/mL interferon for the indicated times. Human cells (HEK293T, HeLa) were stimulated with human IFN-β; MEF cells were stimulated with mouse IFN-α. **B**,**E**,**H**, qPCR analysis of ISGs in HEK293T, HeLa and MEF cells pretreated with 10 µM UA and stimulated with 10ng/mL interferon for the indicated times. Human cells: *ISG15*, *IFIT1*, *OAS1*; MEF: *Isg15*, *Ifit1*, *Oas1*. **C**,**F**,**I**, Densitometric quantification of phosphorylated proteins normalized to the corresponding total proteins in **A**,**D**,**G**, plotted for the indicated times. Statistical significance is indicated: ns = no significance, \**P* < 0.05, \*\**P* < 0.01, \*\*\**P* < 0.001.

UA is a microbiota-derived metabolite associated with mitochondrial quality control ^19,20^. Previous studies have placed UA in pathways that initiate autophagy and execute mitophagy, with ULK1 acting at the initiation step, while Parkin (PRKN) promoting the clearance of damaged mitochondria via a ubiquitin-dependent process ^19,25,26^. Given the established role of mitochondrial homeostasis and selective autophagy in modulating innate immune responses ^27–29^, we next investigated whether the inhibitory effect of UA on IFN-I signaling requires these canonical autophagy components. To address this, we evaluated the effect of UA in *ULK1*-deficient human and murine cells, together with autophagy-deficient cell lines such as *ATG5*-knockout (KO) and *Prkn-*knockout (KO) cell lines (Supplemental Figure 1, D–G). Notably, in these autophagy-deficient cells, UA pretreatment consistently suppressed IFN-I induced phosphorylation of JAK1 and TYK2 without altering total protein levels, with quantification by densitometric scanning of immunoblot bands normalized to the corresponding total protein (Figure 3, A–H), indicating that the inhibitory function of UA is independent of core autophagy machinery.

**Figure 3.**
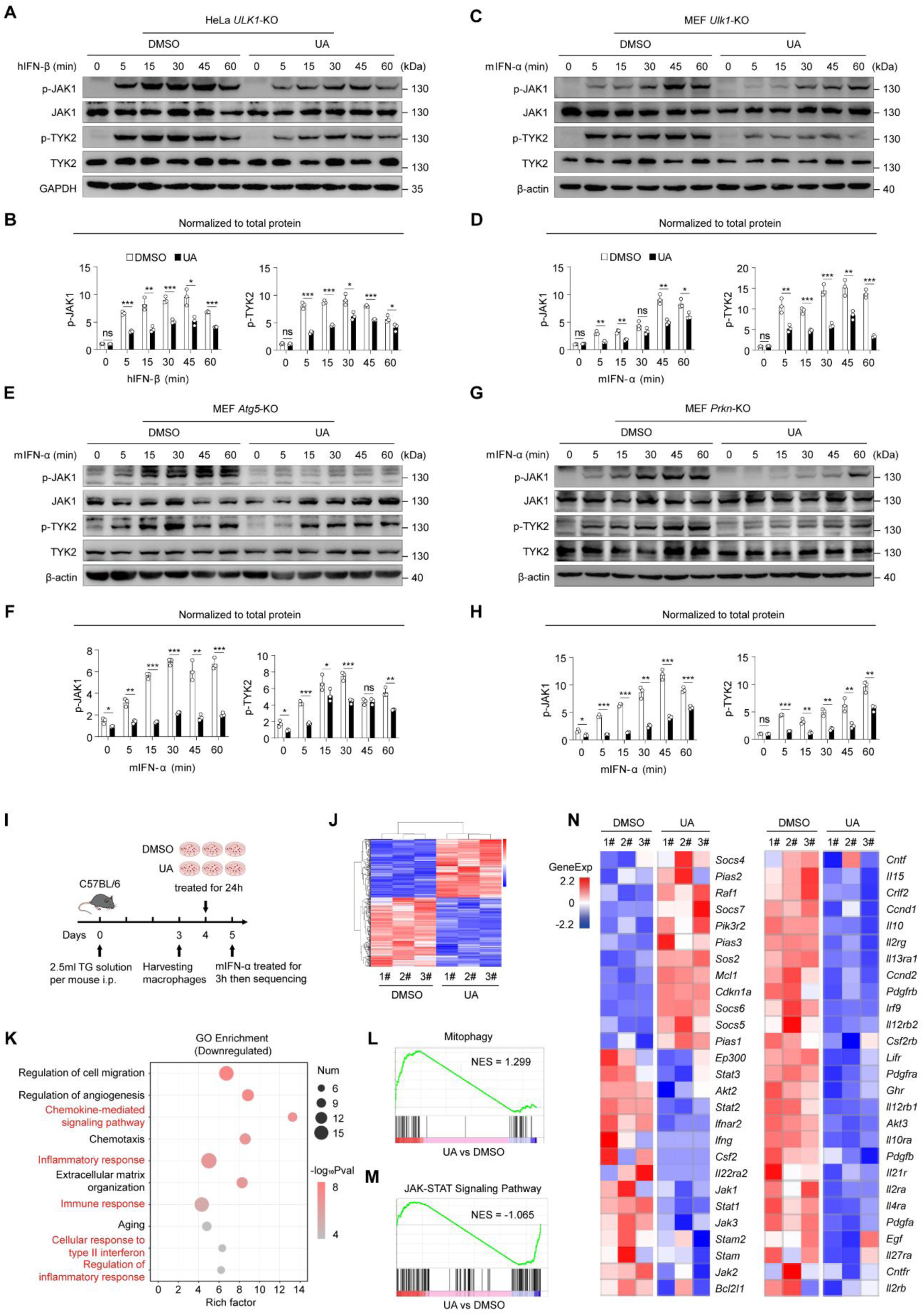
UA limits IFN-I signaling independently of autophagy and represses immune gene-expression programs. **A**,**C**,**E**,**G**, Immunoblot analysis in autophagy-pathway-deficient backgrounds. HeLa *ULK1*-KO, MEF *Ulk1*-KO, MEF *Atg5*-KO and MEF *Prkn*-KO were pretreated with 10 µM UA and then stimulated with 10ng/mL interferon for the indicated times. Human lines were stimulated with human IFN-β; murine lines with mouse IFN-α. **B**,**D**,**F**,**H**, Densitometric quantification of phosphorylated proteins normalized to the corresponding total proteins in **A**,**C**,**E**,**G**, plotted for the indicated times. **I**, Schematic of the RNA-seq design in primary peritoneal macrophages. Cells were exposed to 20 µM UA for 24 h, briefly challenged with 50 ng/mL mouse IFN-α as indicated, and processed for RNA-seq. **J**, Unsupervised sample separation showing distinct transcriptional profiles for UA versus DMSO groups. **K**, Gene Ontology enrichment for downregulated genes. **L**,**M**, GSEA indicating positive enrichment of mitophagy pathway and negative enrichment of the JAK-STAT signaling pathway. **N**, Heat maps of transcripts related to the JAK–STAT signaling pathway. Statistical significance is indicated: ns = no significance, \**P* < 0.05, \*\**P* < 0.01, \*\*\**P* < 0.001.

To elucidate the molecular mechanism of this autophagy-independent suppression, we performed RNA sequencing on primary peritoneal macrophages treated with UA or control and stimulated with IFN-α (Figure 3I). Differential expression analysis revealed a distinct UA transcriptional signature (Figure 3J and Supplemental Figure 1H). Gene set enrichment analysis showed positive enrichment of mitophagy and autophagy programs, consistent with known UA biology (Figure 3L and Supplemental Figure 1, I and J), together with negative enrichment of immune pathways, including the JAK and STAT signaling pathway and cytokine-receptor interaction modules (Figure 3, K and M and Supplemental Figure 1, K–N). The reduced expression of core interferon-stimulated genes and signaling nodes in UA-treated macrophages, spanning the JAK–STAT axes, showed in Heat maps (Figure 3N).

Collectively, these results demonstrate that UA suppresses IFN-I signaling by inhibiting the phosphorylation and activation of JAK1 and TYK2 kinases. This upstream targeting mechanism suggest that UA may act at upstream of the kinases independent of the autophagy processes.

### UA broadly suppresses JAK–STAT signaling downstream of multiple cytokines

Having established UA as a potent inhibitor of IFN-I signaling through JAK–STAT suppression, we next asked whether this activity also applies to other cytokines that signal through the same cascade. Given the central role of JAK kinases in multiple cytokine pathways and their well-documented therapeutic relevance ^10,16,17^, we selected IFN-γ and IL-6 as representative cytokines for further investigation.

For IFN-γ signaling—which critically regulates immune cell functions—we evaluated UA activity in human THP-1 cells and primary mouse peritoneal macrophages, with complementary experiments in HEK293T and MEF cells. UA pretreatment consistently suppressed IFN-γ–induced phosphorylation of JAK1, JAK2, STAT1, and STAT3 across these systems without affecting total protein levels (Figure 4, A, B, D and E and Supplemental Figure 2, A and B). Correspondingly, UA attenuated IFN-γ–driven transcriptional activation, reducing expression of *CXCL9* and *CXCL10* in THP-1 cells and suppressing *Cxcl9* and *Cxcl10* in primary macrophages (Figure 4, C and F).

**Figure 4.**
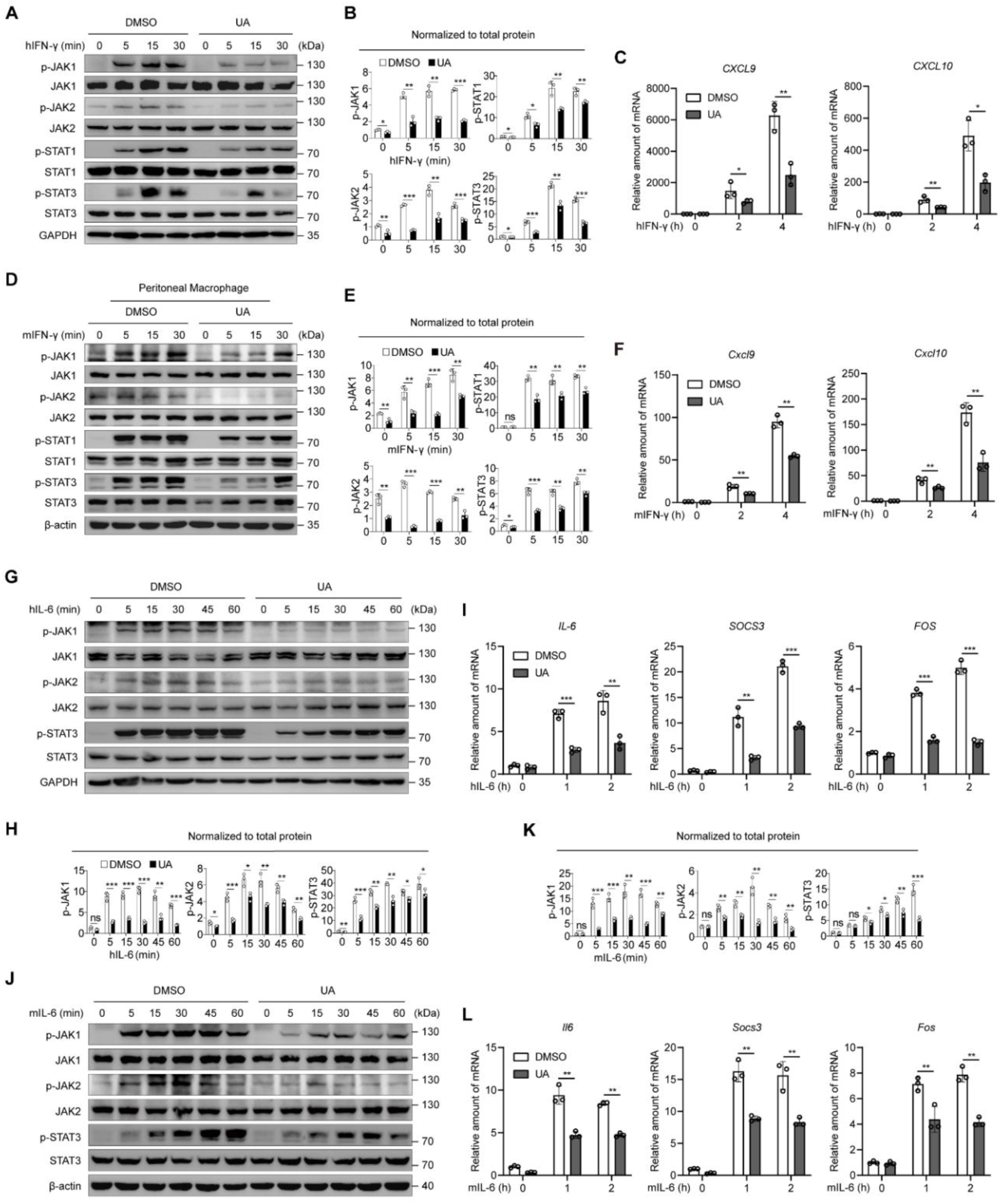
UA suppresses IFN-γ and IL-6 induced signaling. **A**,**D**, Immunoblot analysis of THP-1 cells and primary peritoneal macrophages pretreated with 10 µM UA for 24h and then stimulated with 10ng/mL IFN-γ for the indicated times. THP-1 cells were stimulated with human IFN-γ; primary peritoneal macrophages with mouse IFN-γ. **B**,**E**, Densitometric quantification of phosphorylated proteins normalized to the corresponding total proteins in **A**,**D**, plotted for the indicated times. **C**,**F**, qPCR analysis of IFN-γ pathway targets following UA pretreatment as in **A**,**D**. THP-1: *CXCL9*, *CXCL10*; primary peritoneal macrophages: *Cxcl9*, *Cxcl10*. **G**,**J**, Immunoblot analysis of HEK293T and MEF cells pretreated with 10 µM UA for 24h and then stimulated with IL-6 for the indicated times. HEK293T were stimulated with human IL-6; MEF cells with mouse IL-6. **H**,**K**, Densitometric quantification of phosphorylated proteins normalized to corresponding total proteins in **G**,**J**, plotted for the indicated times. **I**,**L**, qPCR analysis of IL-6 pathway targets following UA pretreatment as in **G**,**J**. HEK293T: *IL6*, *SOCS3*, *FOS*; MEF: *Il6*, *Socs3*, *Fos*. Statistical significance is indicated: ns = no significance, \**P* < 0.05, \*\**P* < 0.01, \*\*\**P* < 0.001.

We similarly examined IL-6 signaling, a pathway with broad inflammatory roles, focusing on HEK293T and MEF cells with validation in primary macrophages. UA pretreatment inhibited IL-6–stimulated phosphorylation of JAK1, JAK2, and STAT3 (Figure 4, G, H, J and K and Supplemental Figure 2C), leading to decreased induction of target genes such as *IL6*, *SOCS3*, and *FOS* in human cells and their orthologs in mouse cells (Figure 4, I and L and Supplemental Figure 2D).

Collectively, these findings demonstrate that UA broadly suppresses JAK–STAT signaling downstream of multiple cytokines across diverse cellular contexts, positioning it as a wide-spectrum modulator of cytokine responses with potential therapeutic implications.

### UA directly binds the JAK1 kinase domain to suppress its catalytic activity

The above findings that UA broadly suppresses JAK–STAT signaling across multiple cytokine pathways led us to investigate whether it directly targets JAK kinases. The convergent inhibition of IFN-α, IFN-γ, and IL-6 signaling, coupled with reduced phosphorylation at receptor-proximal steps, suggested early intervention within the JAK module. Cellular thermal shift assays (CETSA) revealed temperature-dependent stabilization of JAK1 and, to a lesser extent, JAK2 in UA-treated lysates, while TYK2 was largely unaffected (Figure 5, A and B). To exclude indirect effects from JAK1–JAK2 associations, we employed drug affinity responsive target stability (DARTS), which showed that UA conferred pronounced protease resistance to JAK1, with minimal protection of JAK2 and TYK2 (Figure 5, C and D). These complementary approaches consistently identified JAK1 as the primary target.

**Figure 5.**
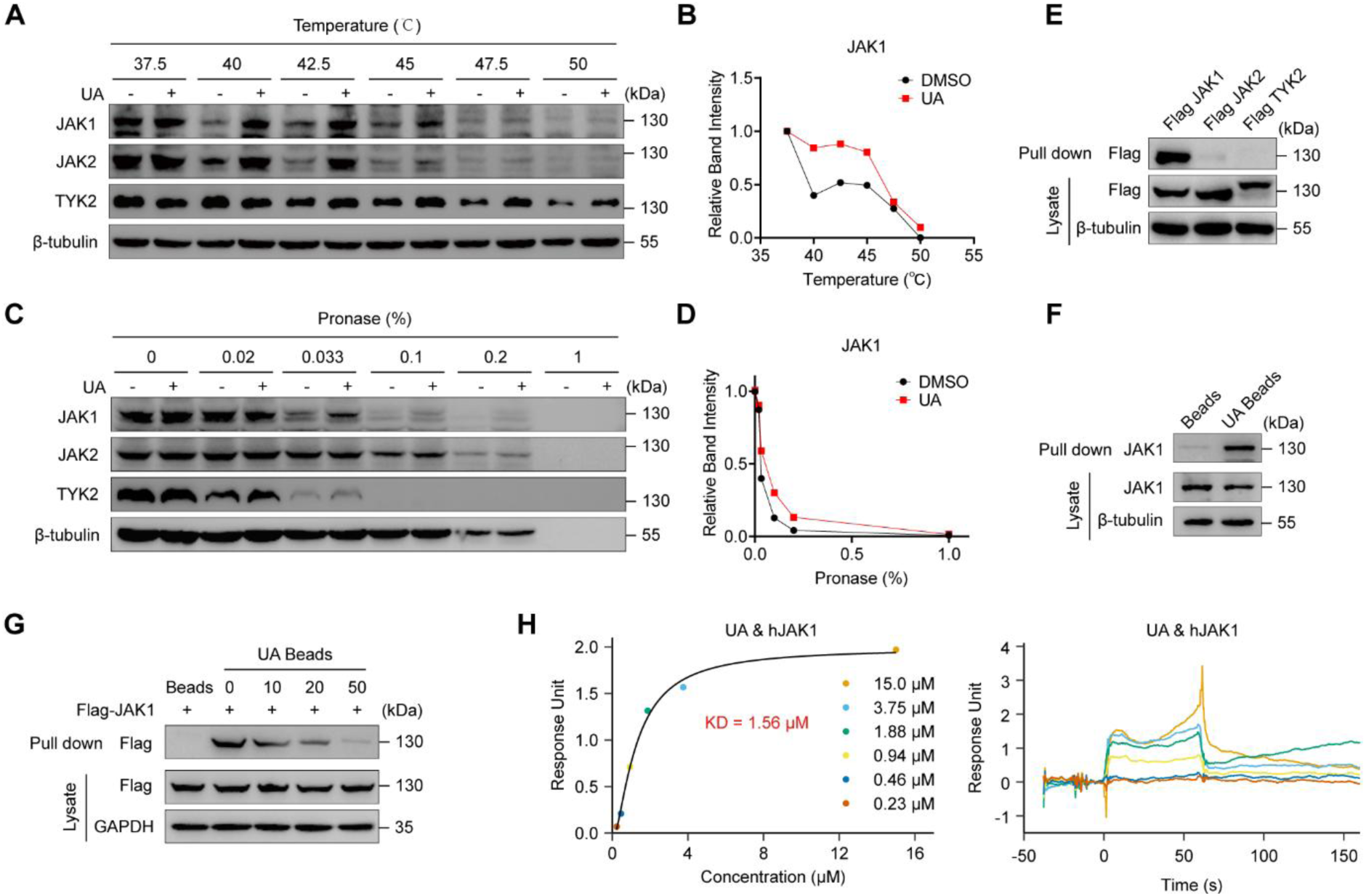
UA selectively engages JAK1 in cells and in vitro. **A**, CETSA of JAK family kinases in UA-treated cell lysates. **B**, Quantification of normalized signal intensities of the JAK1 protein across the indicated temperatures in A. **C**, DARTS assay of JAK family kinases under increasing pronase concentrations. **D**, Quantification of normalized signal intensities of the JAK1 protein across the indicated pronase concentrations in C. **E**, UA-beads pull-down of lysates overexpressing Flag-JAK1, Flag-JAK2 and Flag-TYK2. **F**, UA-beads pull-down of endogenous JAK1 from native lysates. **G**, Competitive binding assay showing that free UA competes with UA-beads for binding to Flag-JAK1. **H**, SPR analysis of UA and hJAK1 protein.

Pull-down assays using UA-conjugated affinity beads further confirmed direct binding, showing efficient capture of Flag-JAK1 but weak or undetectable binding to Flag-JAK2 and Flag-TYK2 (Figure 5E). Endogenous JAK1 was also specifically enriched by UA beads (Figure 5F). Competition experiments demonstrated dose-dependent reduction in JAK1 pulldown by free UA (Figure 5G), and surface plasmon resonance (SPR) confirmed direct interaction between UA and recombinant JAK1 with a KD of 1.56 µM (Figure 5H).

Mechanistically, and consistent with a direct effect on JAK1 signaling output, UA decreased the association between JAK1 and STAT1 following cytokine stimulation (Figure 6, D and E), whereas JAK1–receptor interactions and receptor dimerization were not detectably altered (Figure 6, A–C). To localize the binding interface, a panel of JAK1 truncation constructs was tested: UA-beads selectively enriched full-length JAK1 and the Kinase 2 domain (JH1) but not the FERM, SH2 or Kinase 1 fragments (JH2) (Figure 6F), indicating that the principal UA-binding site resides within the Kinase 2 domain (JH1). Then we performed molecular docking on the hydrophobic pocket of the JH1 domain and found that UA is likely to interact with the three amino acid residues L959, P960, and G1020 in this pocket. (Figure 6G). We also generated a series of JAK1 mutants with single ((L959A,P960A,G1020A), double (L959A/P960A), and triple (L959A/P960A/G1020A) mutants of the predicted pocket residues in the JH1 domain. The result showed that single mutations partially weakened binding, the double mutant exhibited a more pronounced reduction; only the triple mutant (3A) completely abrogated the capture of JAK1, reducing it to nearly undetectable levels (Figure 6H). This stepwise disruption of binding conclusively identifies this pocket as the critical site for the direct interaction with UA. Functional assessment revealed that UA inhibited JAK1 catalytic activity: *in vitro* kinase assays showed diminished JAK1 autophosphorylation and reduced phosphorylation of its substrate STAT1 in the presence of UA, indicating inhibition of both auto- and substrate-directed kinase activity. (Figure 6, I and J). To further determine whether JAK1 is required for the cellular inhibitory activity of UA, we generated CRISPR/Cas9-mediated *JAK1*-knockout (KO) cells. In sgNC control cells, UA markedly reduced hIFN-β-induced phosphorylation of JAK1, TYK2, STAT1, STAT2 and STAT3, accompanied by decreased induction of the canonical ISGs *IFIT1* and *OAS1*. In contrast, JAK1 disruption substantially blunted hIFN-β-induced JAK–STAT activation and ISG transcription, and UA treatment did not confer further detectable inhibition in sgJAK1 cells (Figure 6, K and L).

**Figure 6.**
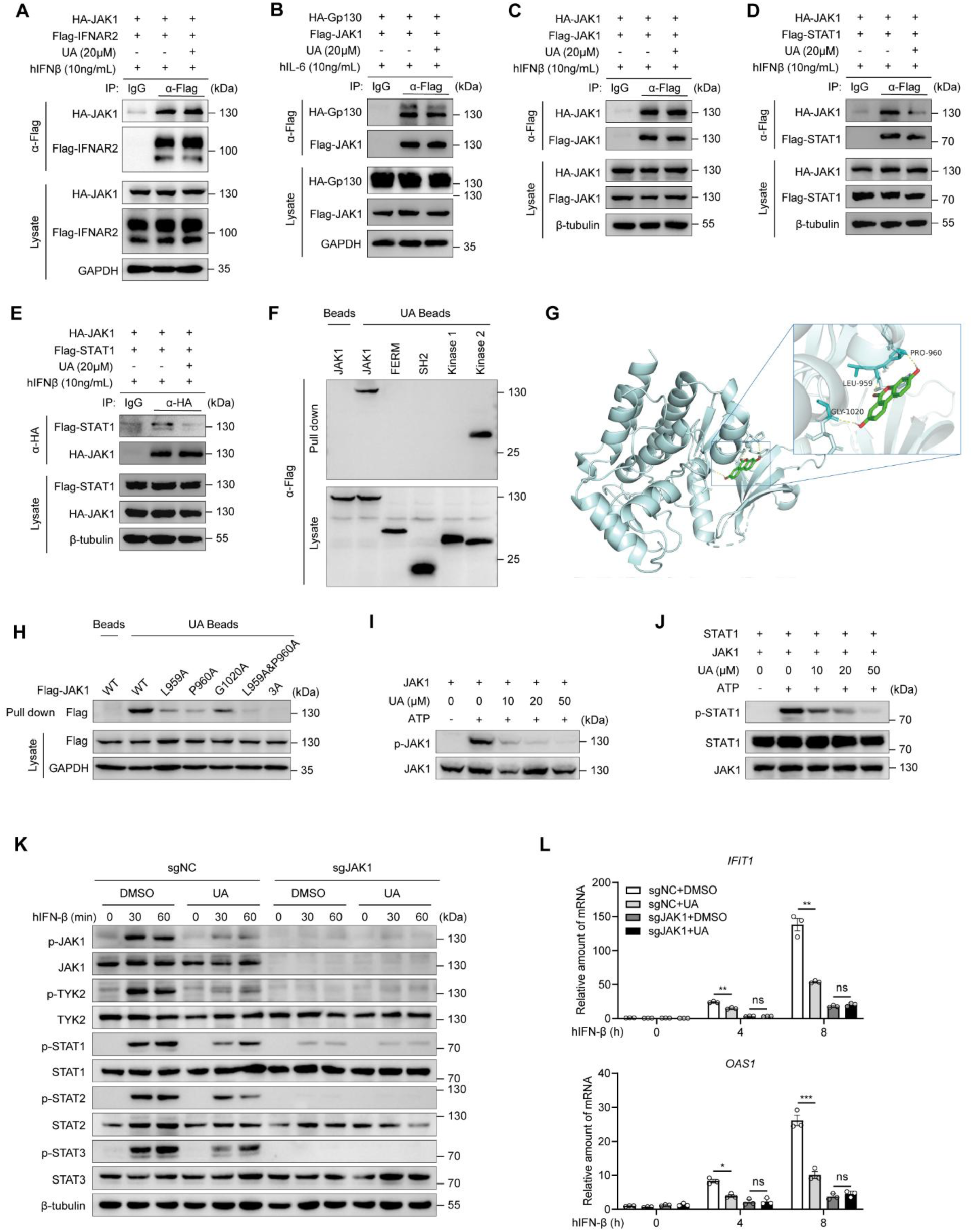
UA targets the JAK1 JH1 domain to suppress catalytic activity and JAK1-dependent STAT signaling. **A**, Immunoprecipitation analysis of the interaction between HA-JAK1 and Flag-IFNAR2 in HEK293T treated with DMSO or UA (20 µM) for 6 h, then stimulated with human IFN-β (10 ng/mL) for 30 min. **B**, Immunoprecipitation analysis of the interaction between HA-Gp130 and Flag-JAK1 in HEK293T treated with DMSO or UA (20 µM) for 6 h, then stimulated with human IL-6 (10 ng/mL) for 30 min. **C**, Immunoprecipitation analysis of the interaction between HA-JAK1 and Flag-JAK1 in HEK293T treated with DMSO or UA (20 µM) for 6 h, then stimulated with human IFN-β (10 ng/mL) for 30 min. **D**,**E**, Immunoprecipitation analysis of the interaction between HA-JAK1 and Flag-STAT1 in HEK293T treated with DMSO or UA (20 µM) for 6 h, then stimulated with human IFN-β (10 ng/mL) for 30 min. **F**, UA-beads pull-down of JAK1 full-length and truncation construct. **G**, Predicted UA-binding pocket within the JAK1 kinase 2 domain identified by molecular docking, highlighting key contact residues. **H**, UA-beads pull-down of JAK1 mutants, showing reduced binding affinity compared with wild-type JAK1. **I**, *In vitro* kinase assay of StrepII-hJAK1 under ATP-containing conditions, showing decreased phosphorylation in the presence of UA. **J**, *In vitro* kinase assay of purified StrepII-hJAK1 with StrepII-STAT1 as a substrate under ATP-containing conditions, showing reduced STAT1 Tyr701 phosphorylation in the presence of UA. **K,** Immunoblot analysis of sgNC control and sgJAK1 HeLa cells pretreated with 10 µM UA for 24 h and then stimulated with 10 ng/mL human IFN-β for the indicated times. **L,** qPCR analysis of ISGs in sgNC control and sgJAK1 HeLa cells pretreated with 10 µM UA for 24 h and stimulated with 10 ng/mL human IFN-β for the indicated times. Human ISGs: *IFIT1*, *OAS1*. Statistical significance is indicated: ns = no significance, \**P* < 0.05, \*\**P* < 0.01, \*\*\**P* < 0.001.

These findings demonstrate that UA directly binds the JAK1 Kinase 2 domain (JH1), impairing its catalytic activity and subsequent STAT1 recruitment. This mechanism provides a structural basis for the broad suppression of UA on cytokine-driven JAK–STAT signaling.

### UA confers protective effects in murine models of JAK–STAT dependent autoimmunity

Building on our mechanistic findings that UA directly targets JAK1 to suppress cytokine signaling, we next evaluated its therapeutic potential *in vivo*. The JAK–STAT pathway serves as a central signaling node for both type I and type II interferon responses, and its dysregulation is implicated in various autoinflammatory diseases. First, we employed the *Trex1*-knockout (KO) model of type I interferonopathy ^30,31^. Loss-of-function mutations in *Trex1* lead to cytosolic DNA accumulation and spontaneous cGAS–STING pathway activation, resulting in persistent IFN-I production and multi-organ inflammation—a phenotype mirroring human Aicardi–Goutières syndrome ^32^. This model directly tests UA’s ability to suppress interferon-driven pathology through JAK1 inhibition. In peritoneal macrophages from *Trex1*-KO mice, transcripts of interferon- and inflammation-associated genes, including the canonical ISG *Oas1* as well as *Cxcl10* and *Il6*, were significantly elevated compared to wild-type controls, and oral UA treatment substantially reduced their expression (Figure 7A). After 14 days of UA administration (Figure 7B), histopathological analysis revealed attenuated inflammatory injury in heart, kidney, and skeletal muscle, with improved tissue architecture and lower histological scores in UA-treated *Trex1*-KO mice (Figure 7, C and D). Concordantly, multi-organ qPCR confirmed decreased expression of *Il6*, *Cxcl10*, *Isg15*, and *Oas1* (Figure 7E), demonstrating UA’s capacity to dampen interferon-driven inflammation in a genetic model of IFN-I dysregulation.

**Figure 7.**
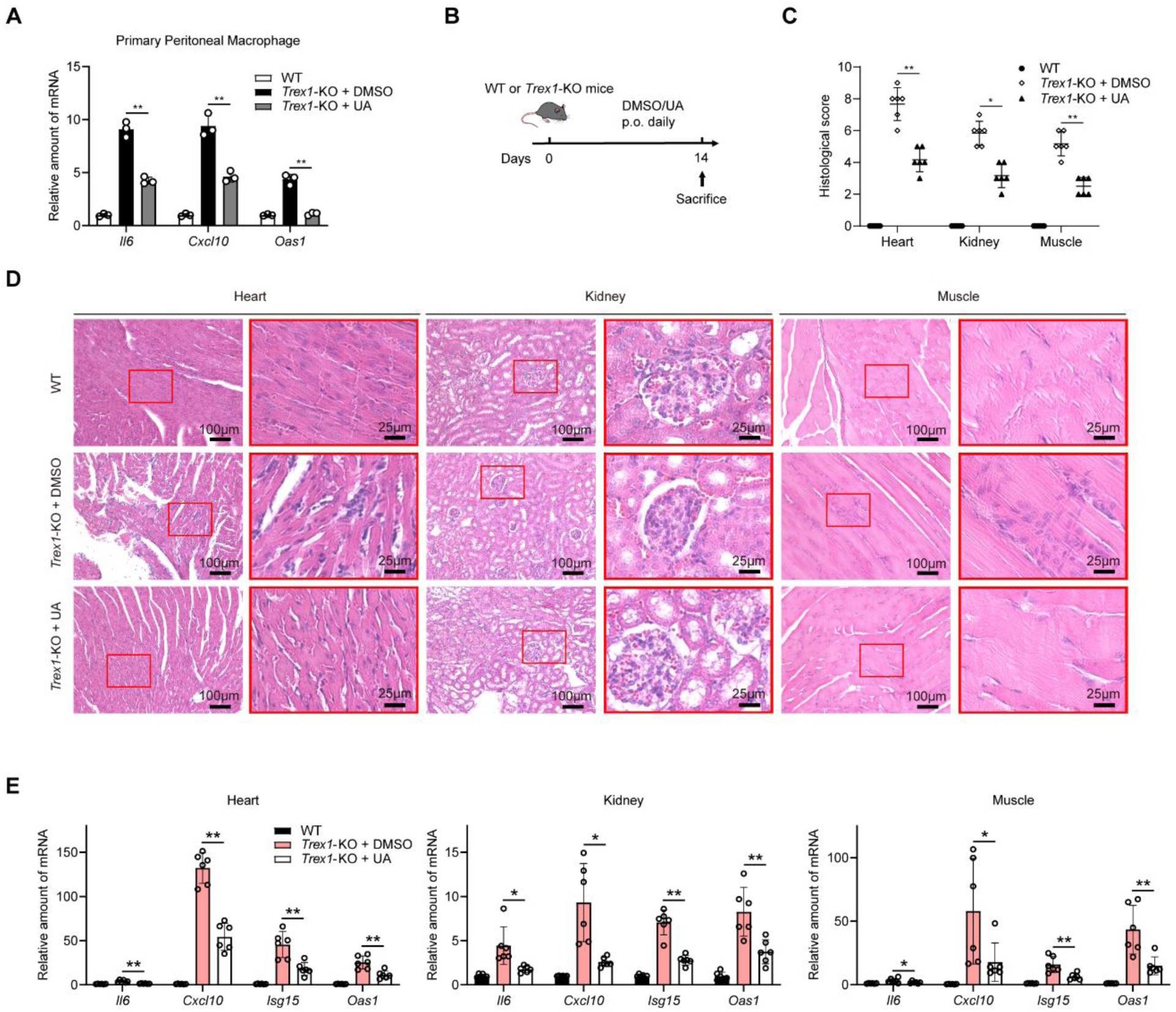
UA alleviates interferon-driven inflammation in *Trex1*-KO mice. **A**, qPCR analysis of peritoneal macrophages from WT and *Trex1*-knockout mice with or without UA treatment. **B**, Schematic of the *in vivo* treatment protocol (n = 6 per group). *Trex1*-KO and WT mice received oral UA(100mg/kg) or DMSO once daily for 14 days before sample collection. **C**, Quantification of histopathological scores for heart, kidney and skeletal muscle from WT and *Trex1*-KO mice with or without UA treatment. **D**, Representative histology of heart, kidney and skeletal muscle from WT and *Trex1*-KO mice with or without UA treatment. **E**, qPCR analysis of ISGs (*Il6*, *Cxcl10*, *Isg15*, *Oas1*) in heart, kidney and muscle from WT and *Trex1*-KO mice with or without UA treatment. Statistical significance is indicated: ns = no significance, \**P* < 0.05, \*\**P* < 0.01, \*\*\**P* < 0.001.

To evaluate UA’s efficacy in a cytokine-rich inflammatory environment, we utilized the imiquimod (IMQ)-induced psoriasis model, where TLR7/8 activation triggers IL-23/IL-17 axis activation and JAK–STAT–dependent cytokine production ^33,34^. During IMQ challenge (Supplemental Figure 3A), UA treatment reduced splenomegaly (Supplemental Figure 3B), alleviated clinical symptoms (erythema, scaling, epidermal thickening), and improved PASI scores (Supplemental Figure 3, C and D). Histology showed diminished epidermal hyperplasia and immune infiltration (Supplemental Figure 3E), while lesional skin qPCR revealed downregulation of JAK–STAT–responsive genes (*Isg15*, *Ifit1*, *Oas1*) and pro-inflammatory mediators (*Tnf*, *Il6*) (Supplemental Figure 3F).

To further assess the therapeutic potential of UA in systemic autoimmunity, we employed a low-frequency, 6-week imiquimod (IMQ)-induced systemic lupus erythematosus (SLE) model in BALB/c mice (Figure 8A) — a model driven by sustained TLR7 stimulation and associated interferon signatures ^35^. Compared with controls, IMQ challenge induced marked splenomegaly (increased spleen size, weight, and index), which was substantially mitigated by UA treatment (Figure 8, B and C). Consistent with attenuation of systemic autoreactivity, UA reduced circulating anti–dsDNA antibody levels measured by ELISA (Figure 8D). Renal function was similarly improved, as evidenced by a significant decrease in the urinary albumin-to-creatinine ratio (uACR) (Figure 8E). Histopathological examination of kidneys revealed that UA alleviated IMQ-induced glomerular and interstitial injury on H&E staining, accompanied by lower histological scores (Figure 8, F and H). In parallel, immunofluorescence staining demonstrated reduced renal IgG deposition in UA-treated mice, supported by semiquantitative analysis of fluorescence intensity (Figure 8, G and I). At the transcriptional level, UA suppressed the IMQ-induced upregulation of interferon/JAK–STAT–responsive genes (e.g., *Isg15*, *Cxcl10*) in both spleen and kidney tissues (Figure 8, J and K).

**Figure 8.**
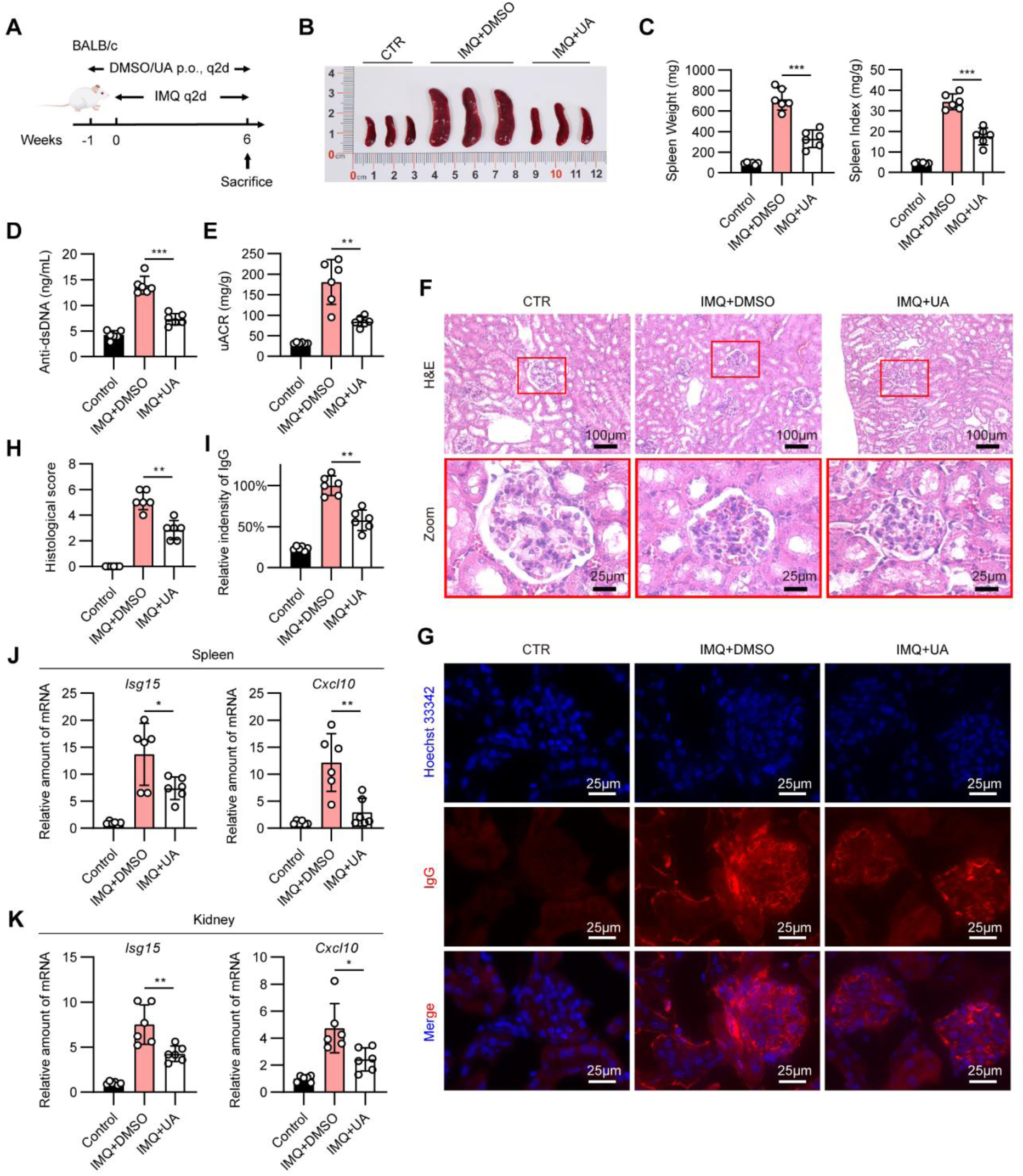
UA mitigates systemic autoimmunity and renal injury in IMQ-induced SLE model. **A**, Schematic of the in vivo induction and treatment protocol in BALB/c mice. Mice received IMQ every 2 days and were administered oral UA or DMSO every 2 days for 6 weeks before sacrifice (n = 6 mice per group). **B**, Representative gross morphology of spleens from control, IMQ+DMSO and IMQ+UA groups. **C**, Quantification of spleen weight and spleen index in control, IMQ+DMSO and IMQ+UA groups. **D**, Plasma anti-dsDNA antibody levels measured by ELISA. **E**, Urinary albumin-to-creatinine ratio (uACR) calculated from ELISA-measured urine albumin and creatinine (n = 6 mice per group). **F**, Representative H&E staining of kidney sections from control, IMQ+DMSO and IMQ+UA groups, with boxed areas shown at higher magnification. **G**, Representative immunofluorescence staining of renal IgG deposition with Hoechst 33342 counterstaining in control, IMQ+DMSO and IMQ+UA groups. **H**, Quantification of renal histopathological scores based on H&E-stained kidney sections. I, Semiquantification of renal IgG deposition based on relative fluorescence intensity. **J**, qPCR analysis of ISGs (*Isg15*, *Cxcl10*) in spleen tissues from control, IMQ+DMSO and IMQ+UA groups. **K**, qPCR analysis of ISGs (Isg15, Cxcl10) in kidney tissues from control, IMQ+DMSO and IMQ+UA groups. Statistical significance is indicated: ns = no significance, \**P* < 0.05, \*\**P* < 0.01, \*\*\**P* < 0.001.

Therefore, UA represents a novel JAK1 inhibitor with significant therapeutic potential. Our findings demonstrate its unique mechanism of action through direct kinase domain binding and validate its efficacy in suppressing pathological inflammation across different autoimmune models. These results support the further development of UA as a promising therapeutic agent for JAK–STAT-driven autoimmune disorders.

## Discussion

Dysregulated type I interferon signaling is the underlying cause of a range of immune pathologies, from monogenic interferonopathies such as Aicardi-Goutières syndrome (AGS) and STING-associated vasculopathy with onset in infancy (SAVI), to systemic autoimmune diseases including systemic lupus erythematosus (SLE) ^36,37^. The hallmark of these conditions is sustained activation of IFN-I pathway, underscoring the therapeutic rationale for targeting this signaling axis through various strategies. Current approaches include the direct blockade of the IFN-I pathway using monoclonal antibodies (e.g., anifrolumab targeting IFNAR), as well as the inhibition of downstream signaling via JAK inhibitors (e.g., tofacitinib, baricitinib) ^16,17,34^. While these targeted biologics and synthetic drugs demonstrate clinical efficacy, their use can be limited by safety concerns and other side effects, such as an increased risk of infections and drug resistance. These limitations underscore the need to develop safer and more tolerable therapeutic agents ^16,17,34^. Research should focus not only on refining selective JAK inhibitors but also on exploring naturally derived or low-cost compounds that can modulate this pathway. Such efforts could expand treatment accessibility while providing novel chemical scaffolds for therapeutic development.

Certain gut microbial metabolites act as indirect modulators of the JAK–STAT pathway, playing a key role in regulating immune homeostasis and controlling inflammation. For example, butyrate, produced by *Faecalibacterium prausnitzii,* suppresses JAK–STAT activation by enhancing the histone acetylation at the suppressor of cytokine signaling 1 (SOCS1) promoter and promoting its expression, which is known as a negative regulator of the JAK–STAT pathway ^5^. Acetate derived from gut microbiota is reported to activate JAK1/STAT3 to drive M1 macrophage polarization ^38^. Urolithin A is a gut microbiota-derived metabolite generated from ellagitannin-rich foods and has been extensively characterized as a mitophagy-activating and broadly anti-inflammatory agent. UA promotes PINK1/Parkin-mediated mitophagy, thereby clearing damaged mitochondria and reducing mitochondrial ROS-dependent NLRP3 inflammasome activation ^24^. In parallel, UA engages energy-sensing pathways (for example, AMPK) and dampens NF-κB signaling, while in the adaptive compartment it promotes regulatory T cell (Treg) expansion and constrains Th1/Th17 differentiation, restoring immune balance across diverse inflammatory settings ^39–41^. Inter-individual capacity to produce UA is strongly influenced by gut microbiome composition ^18^; low UA producers in human cohorts display higher systemic inflammatory markers such as C-reactive protein (CRP) ^42^. These observations suggest that UA insufficiency may contribute to chronic inflammation in a subset of individuals.

Although UA is well established to exert anti-inflammatory effects, there has been no direct evidence that it interacts with key cytokine signaling components, such as the JAK–STAT pathway, or that it regulates cytokine-driven immune responses. Here we identify UA as a previously unrecognized regulator of cytokine-driven JAK–STAT signaling. Across multiple cell types, UA uniformly suppressed JAK–STAT activation triggered by type I IFNs, type II IFN and IL-6 and curtailed early phosphorylation of JAK1, JAK2 and TYK2, consistent with an action at or immediately upstream of the JAK module rather than at the level of STATs or distal transcriptional events. Transcriptomic profiling of primary macrophages corroborated these findings, revealing broad attenuation of interferon-stimulated gene programs and JAK–STAT pathway signatures upon UA exposure. Importantly, genetic disruption of *ULK1*, *Atg5* or *Prkn* did not relieve UA-mediated inhibition of cytokine signaling, indicating that this effect is independent of the mitophagy machinery that has dominated the conceptualization of UA biology. This is mechanistically notable because most anti-inflammatory effects of UA have been attributed to enhanced the clearance of damaged mitochondria and inflammasome activators. By showing that UA still dampens IFN- and IL-6–induced JAK–STAT activation in cells lacking core autophagy components, our data delineate a parallel signaling arm in which UA modulates cytokine receptor–proximal events without overtly relying on autophagic flux. To test whether UA can target at the kinase level, UA beads assay was performed, and it was demonstrated that UA can bind directly to JAK1, rather than JAK2 or TYK2. This interaction was also supported by thermal shift assays and surface plasmon resonance (SPR). These results suggest that UA may be a natural JAK1 inhibitor.

Pull-down assays confirmed the direct interaction between UA and JAK1. Site-directed mutagenesis mapped the critical binding determinants to a three-residue cluster comprising L959, P960 and G1020. Structurally, L959 and P960 reside in the hinge region of the ATP-binding pocket, while G1020 is located within the activation loop of the JH1 domain. This places the UA-binding site within the same structural compartment as ATP-competitive JAK inhibitors, including clinically used agents such as tofacitinib and baricitinib, which typically occupy the JH1 ATP-binding cleft and form key hydrogen bonds with hinge residues such as E957 and L959 in JAK1 or the equivalent residues in other JAKs ^43,44^. In contrast to typical ATP-competitive scaffolds, however, UA engages the canonical hinge residue L959 together with the adjacent hinge residue P960, and the functional sensitivity of G1020 suggests that the activation loop also contributes to binding. Whether this extended hinge–activation loop interface translates into a distinct selectivity profile compared with existing JAK1 inhibitors will require future structural and pharmacological comparison. Given that selective JAK1 inhibition preferentially attenuates autoimmune-relevant cytokine signaling (for example, IFNs, IL-6 and IL-23) while relatively sparing JAK2-dependent hematopoiesis ^45^, our findings support UA as a promising starting point for the development of metabolite-inspired JAK1 modulators for autoimmune diseases. The CRISPR/Cas9-mediated *JAK1*-KO cell line also provided important genetic support for the proposed mechanism. UA effectively suppressed IFN-β-induced JAK–STAT phosphorylation and ISG expression in sgNC control cells, whereas this inhibitory effect was largely lost in *JAK1*-KO cells, in which IFN-I signaling was already markedly impaired. These results link the direct biochemical interaction between UA and JAK1 to its cellular activity and support JAK1 as a functionally relevant target of UA. However, because JAK1 deletion compromises the integrity of receptor-proximal IFN-I signaling, future rescue experiments with wild-type JAK1 or UA-binding-deficient mutants, particularly the L959A/P960A/G1020A triple mutant, will help determine whether this JH1 pocket is required for UA-mediated inhibition in living cells.

To examine whether UA has therapeutic potential in autoimmune diseases, *Trex1*-KO mice was used, which is a type of mouse model with excessive type I interferons ^30,32^. UA treatment can significantly reduce the disease progression and severity. We also tested another disease model, using IMQ-induced psoriasis and SLE models ^33^, UA also have beneficial effects. Taken together, our findings identify JAK1 as a central signaling node in cytokine-driven autoimmunity that can be modulated by the microbiota-derived metabolite UA.

UA exhibits a favorable preclinical and clinical safety profile, supporting its potential for therapeutic application. In 28- and 90-day repeated-dose oral toxicity studies in rats, synthetic UA administered at dietary concentrations of up to 5% exhibited no genotoxicity, target-organ toxicity, or clinically significant alterations in clinical parameters ^46^. The highest dose tested was established as the no-observed-adverse-effect level. In human trials, multiple randomized, placebo-controlled studies involving middle-aged and older adults (including NCT03283462, NCT03464500, and NCT05735886) have consistently demonstrated that once-daily UA supplementation at doses of 250–1,000 mg for durations ranging from 4 weeks to 4 months is safe and well tolerated ^20–22^. These studies also reported improvements in biomarkers related to mitochondrial function, muscle performance, systemic inflammation (e.g., CRP and specific ceramides), and immune aging, with no increase in adverse events. It is clinically imperative to assess the therapeutic effects of UA on chronic inflammatory and autoimmune conditions. Strategies to enhance UA levels—either through dietary intervention, microbiota modulation, or direct supplementation—warrant further investigation as potential approaches for immune regulation.

Beyond JAK1, UA is also reported to activate mitophagy, targeting ERK1/2, AhR, and GLS1 ^41,47,48^. Future research must systematically identify potential additional targets and, crucially, determine which specific interaction is most critical for its core anti-inflammatory activity. Furthermore, a key strategic question is whether this multi-target profile can be leveraged to achieve synergistic benefits—concurrently combating senescence, suppressing inflammation, and enhancing muscle performance and anti-tumor immunity.

In summary, we demonstrated for the first time that UA directly targets JAK1 and inhibits the activation of JAK–STAT pathway. It has translational implications in chronic inflammatory and autoimmune diseases.

## Methods

### Sex as a biological variable

Autoimmune diseases, including systemic lupus erythematosus and psoriasis, show a marked female predominance in humans. Therefore, all *in vivo* experiments were performed in female mice to maximize clinical relevance and to minimize variability attributable to sex-specific differences in immune responses. Sex was not a biological variable in cell-based experiments. The findings may not fully capture sex-dependent differences, and additional studies in male mice will be necessary to evaluate the generalizability of these effects across sexes.

### Cell lines

The following cell lines were maintained in Dulbecco’s Modified Eagle Medium (DMEM, GIBCO): HEK293T, HeLa Wild-type (WT), HeLa *ULK1* knockout (KO), immortalized mouse embryonic fibroblast (MEF) WT, MEF *Ulk1* KO, MEF *Atg5* KO, MEF *Prkn* KO, L929-ISRE, and 2fTGH-ISRE cells.

Primary peritoneal macrophages were cultured in RPMI 1640 (GIBCO). All media used were supplemented with 10% (v/v) fetal bovine serum (FBS), 100 U/ml penicillin, and 0.1 mg/ml streptomycin.

### Plasmids and transfection

Plasmids used in this study, including pRK-HA-JAK1, pRK-Flag-JAK1(and its truncated mutants), pRK-Flag-TYK2, pCS2-StrepⅡ-Flag-JAK1 and pRK-Flag-STAT1 were generated by standard molecular cloning or site-directed mutagenesis (for mutants) as previously described, with all constructs verified by sequencing. pENTER-Flag-JAK2 and pCMV-Flag-IFNAR2 were obtained from WZ Biosciences and MiaoLing biology, respectively. pCMV-HA-Gp130 was a gift from Dr. Hong-Bing Shu (Wuhan University, China).

For transient expression, HEK293T cells were transfected with polyethylenimine (PEI; Polysciences) following the manufacturer’s protocol. Cells were harvested 24 to 48 hours post-transfection for subsequent analysis.

### Cell viability measurement

Cell viability was assessed using the Cell Counting Kit-8 (AQlabtech). In detail, cells were seeded in 96-well plates and allowed to adhere overnight. The following day, the cells were treated with UA (Selleck, cat#S5312) at 10, 20, 30, 50 and 150 µM or vehicle control(the same amount of DMSO) for 24 h. Thereafter, 10 µL of CCK-8 reagent was added to each well, and the plates were incubated for 1–2 h at 37 °C before the absorbance at 450 nm was measured. Each condition was assayed in at least technical triplicate.

### RNA extraction and quantitative real-time PCR( qRT–PCR) analysis

Total RNA was extracted from cells using TRIzol reagent (Yeasen). Complementary DNA (cDNA) was then synthesized from equal amounts of RNA using a reverse-transcription kit (Yeasen). Quantitative PCR using SYBR Green master mix (Yeasen) on a TIANLONG real-time PCR system. Gene expression was normalized to *Gapdh* or *Actb*, and relative quantification was performed using the 2^–ΔΔCt method. All primer sequences used are provided in Supplemental Table 2.

### Immunoblotting

Cells were lysed on ice using RIPA buffer (APPLYGEN) containing protease and phosphatase inhibitors. Lysates were cleared by centrifugation (12,000 rpm, 10 min, 4 °C). Protein concentrations were determined by a BCA assay (APPLYGEN), and equal amounts of protein were separated by SDS-PAGE. Electrophoresis was performed at 65 V for 40 min throughout the stacking gel, followed by 120 V for 40 min. Proteins were then transferred onto nitrocellulose (NC) membranes at 350 mA for 65 min. The membranes were blocked with 6% (w/v) non-fat milk in TBST at room temperature for 1 h, and subsequently incubated with primary antibodies (diluted in TBST with 5% BSA or as recommended) overnight at 4 °C. After washing with TBST, membranes were incubated with HRP-conjugated secondary antibodies for 1 h at room temperature. Blots were visualized by enhanced chemiluminescence (ECL) substrate and imaged with Image Quant LAS 500. Band intensities were quantified using ImageJ software and normalized to the corresponding total protein or loading control. All antibodies are detailed in Supplemental Table 1.

### Co-immunoprecipitation

HEK293T cells were transfected and cultured for 24 h prior to harvesting. Cells were lysed in ice-cold co-IP buffer (for 1 h at 4 °C) supplemented with a protease inhibitor cocktail (Roche) and 1mM PMSF (Sigma-Aldrich). Cell lysates were clarified by centrifugation, and the supernatants were incubated with the indicated primary antibodies for immunoprecipitation. Subsequently, protein A agarose beads (GE Healthcare) were added to the lysate-antibody mixture, and incubation continued for additional 3–4 h at 4 °C with rotation. The beads were then collected and washed five times with lysis buffer to remove non-specifically bound protein, resuspended in SDS loading buffer, and boiled at 95 °C for 10 min. Both the input lysates and co-immunoprecipitated samples were resolved by SDS-PAGE and subjected to western blot analysis with the relevant antibodies.

### Luciferase reporter assay

To assess the effect of UA on IFN signaling, 2fTGH and L929 cell lines stably expressing an interferon-stimulated response element (ISRE)-driven firefly luciferase reporter were seeded in 96-well plated. After 24 hours of pretreatment with UA, cells were stimulated with 10ng/mL human IFN-β (SinoBiological) or 10ng/mL mouse IFN-α (Novoprotein) for 12 h. Luciferase activity was then assessed using the Dual-Luciferase Reporter Assay System (TransGen) following the manufacturer’s protocol.

### Cellular thermal shift assay (CETSA)

To evaluate the binding between UA and JAK kinases, CETSA was performed ^49^. HEK293T cells (∼80% confluent) were treated with 10 µM UA or vehicle for 24 h. Cells were then harvested, washed with cold PBS and resuspended in PBS supplemented with protease inhibitor cocktail and PMSF. Aliquots of the cell suspension were heated at the indicated temperatures (37.5 °C, 40 °C, 42.5 °C, 44 °C, 50 °C) for 3 min using a thermocycler, followed by three freeze-thaw cycles (liquid nitrogen to room temperature) for lysis. After centrifugation, the soluble fractions were subjected to immunoblotting using antibodies against JAK1, JAK2, and TYK2.

### Drug affinity-responsive target stability (DARTS)

For DARTS Assay ^50^, HEK293T cells were grown in 10 cm dishes and treated for 24 h with 10 µM UA or vehicle for 24 h. Cells were lysed in NP-40 lysis buffer with protease inhibitors and PMSF. After centrifugation, protein concentration was determined by BCA. Lysate aliquots (250µg each) were treated with a pronase (MCE) gradient (0, 0.05, 0.083, 0.25, 0.5, and 2.5 µg per tube) for 30 min at 37 °C. Reactions were terminated by adding a protease inhibitor cocktail and placing on ice for 10 min. Samples were then prepared in Loading buffer and analyzed by immunoblotting with antibodies against JAK1, JAK2, and TYK2.

### UA-beads pull-down and competition assay

UA was covalently immobilized onto NHS-activated magnetic beads (Yeasen) following an established protocol ^47^. For the pull-down assay, lysates from HEK293T cells overexpressing Flag-JAK1, Flag-JAK2, or Flag-TYK2 were incubated with UA-conjugated magnetic beads or control magnetic beads for 4 h at 4 °C. The bead-bound proteins were subsequently eluted and analyzed by immunoblotting. For the competition assay, cell lysates were incubated with control magnetic beads or UA-conjugated beads, alongside UA at 10, 20, and 50 μM doses.

### Surface plasmon resonance (SPR)

StrepII-tagged human JAK1 (StrepII-hJAK1) was transiently overexpressed in HEK293T cells. The protein was purified from clarified lysates using Strep(II)Sep Agarose Purification Resin 4FF (Yeasen) according to the manufacturer’s protocol. The purified protein was concentrated to > 0.5 mg/mL via ultrafiltration and desalted into running buffer with a Cytiva desalting column. All SPR experiments were performed on a Biacore 8K instrument (Cytiva) at 25 °C. HBS-P+ running buffer (10 mM HEPES pH 7.4, 150 mM NaCl, 0.05% Tween-20) was used throughout. Purified JAK1 was immobilized on a CM5 sensor chip (Cytiva) via standard amine-coupling. Compound UA was injected at a series of concentrations in running buffer with a constant flow rate at 30 µL/min. The equilibrium dissociation constant KD value was determined using the Biacore 8K Evaluation Software.

### Molecular docking

The structure of JAK1 kinase 2 domain (JH1) (PDB ID: 6N77) was preprocessed in PyMOL by removing waters/ions and alternates, adding hydrogens, and saving the domain as the receptor. Urolithin A was prepared in AutoDockTools by adding hydrogens, defining rotatable bonds and assigning Gasteiger charges. Both the receptor and ligand were exported as PDBQT files. Molecular docking was performed using AutoDock Vina, with the grid centered on the kinase 2 (JH1) pocket. The predicted binding free energies (ΔG) were recorded, and the lowest energy was selected for analysis. The top-ranked binding poses were visualized and structurally annotated using PyMOL.

### In vitro kinase assay

StrepII-tagged human JAK1 (StrepII-hJAK1) was purified as described in the previous section. Kinase reactions were carried out by incubating purified StrepII-hJAK1 with specified concentrations of UA in kinase assay buffer (10 mM HEPES pH 7.4, 150 mM NaCl, 8 mM MgCl₂, 2 mM MnCl₂, 1 mM DTT, 2mM ATP) in a total volume of 20-25 µL at 30 °C for 40 min. The reactions were terminated by adding 5 × SDS-PAGE loading buffer and boiling for 5 min. Samples were resolved by SDS-PAGE and phospho-JAK1 levels was detected by immunoblotting using a specific anti-phospho-JAK1 antibody. Following membrane stripping, total JAK1 level was assessed using an anti-JAK1 antibody. Both antibodies are detailed in Supplemental Table 1.

### CRISPR/Cas9 knockout

HeLa *JAK1*-KO cells were generated using the lentiCRISPR-V2 system. Double-stranded oligonucleotides corresponding to the sgRNA target sequence against human JAK1 were annealed and cloned into the BsmBI-digested lentiCRISPR-V2 vector. The sgRNA sequence was sgJAK1: 5′-GCTGCCTCGAAGAAGGCCTG-3′. A non-targeting sgRNA was used as the negative control. Lentiviral particles were produced by cotransfecting HEK293T cells with lentiCRISPR-V2-sgRNA, psPAX2 and pMD2.G packaging plasmids using polyethyleneimine. Viral supernatants were collected after transfection and used to transduce HeLa cells in the presence of polybrene. Twenty-four hours after transduction, the medium was replaced with fresh complete medium. After an additional 24 h, infected cells were selected with puromycin at 2 μg/mL for 4 d. Surviving sgJAK1-1-transduced cells were subjected to single-cell cloning by limiting dilution, and individual clones were expanded JAK1 knockout efficiency was validated by immunoblot analysis, and one validated sgJAK1-1-derived monoclonal cell line was used for subsequent experiments. The sgNC control cells were generated and selected in parallel under the same conditions.

### Isolation of primary peritoneal macrophages

Primary peritoneal macrophages were elicited by intraperitoneal injection of 2.5 ml of 3% (w/v) thioglycollate medium to C57BL/6J mice. Three days post-injection, peritoneal exudate cells were harvested by lavage with ice-cold PBS. The cells were plated in RPMI-1640 medium supplemented with 10% (v/v) FBS. After a 2-hour incubation, the non-adherent cells were removed by gentle washing. After equilibration in fresh complete medium, adherent cells were treated with 20 µM UA for 24 h. Following UA exposure, cells were processed for downstream assays.

### RNA-seq analysis

Primary peritoneal macrophages were isolated as described above. Cells were treated with 20 µM UA for 24 h, followed by stimulation with mouse IFN-α (50 ng/mL) for 4 h. Cells were then harvested, and total RNA was extracted using TRIzol reagent. Library construction, high-throughput sequencing, and primary data processing (including quality control, adapter trimming and read mapping) were conducted by NovelBio (Shanghai, China) according to their standard protocols. Differential gene expression analysis was performed with a significance threshold set at |log₂ fold-change| > 1.5 and an adjusted *P*-value < 0.05. Functional enrichment analyses, including Gene Ontology (GO) and Gene Set Enrichment Analysis (GSEA), were carried out using the clusterProfiler with gene sets obtained from the MSigDB database. Results were visualized using standard plotting tools.

### Animal experiments

Both C57BL/6 and BALB/c mice were purchased from Beijing Vital River Laboratory Animal Technology Co., Ltd. (Beijing, China). The *Trex1*-KO mice were kindly provided by Dr. Tomas Lindahl (The Francis Crick Institute, United Kingdom). All mice were maintained under specific pathogen-free (SPF) conditions.

General animal procedures. Female mice were randomized to treatment groups as indicated. UA was administered by oral gavage at 100 mg/kg formulated in vehicle (5% DMSO, 40% PEG300, 5% Tween-80, and 50% saline) according to the schedules described below. At the indicated endpoints, mice were euthanized and tissues were collected for histology, immunofluorescence, ELISA, and RNA extraction for RT–qPCR, as specified for each model. IFN-α response model. C57BL/6 mice were dosed with UA (100 mg/kg, oral gavage) daily for 7 days, followed by a single tail-vein injection of mouse IFN-α (100 ng/g body weight). Four hours after IFN-α injection, organs were harvested, snap-frozen in liquid nitrogen, and stored at –80 °C for RNA extraction and RT–qPCR analysis of ISG expression.

### *Trex1*-KO autoimmune model

*Trex1*-knockout mice (C57BL/6 background) were genotyped by PCR and used at 4–5 weeks of age. Mice were randomized into WT, KO + vehicle, and KO + UA groups and treated by oral gavage with UA (100 mg/kg) or vehicle daily for 14 days. Hearts, kidneys, and skeletal muscles were collected for histology ^51^ and qPCR analysis.

IMQ-induced psoriasis model. Female BALB/c mice were treated by oral gavage with UA (100 mg/kg) or vehicle once daily starting 7 days before IMQ challenge and throughout the experiment. Psoriasis-like inflammation was induced by daily topical application of 62.5 mg of 5% imiquimod cream (Mingxin Pharmaceutical) to shaved dorsal skin for 5 consecutive days. Disease severity and tissue collection were performed at the indicated time points as described ^52^.

IMQ-induced SLE model. Female BALB/c mice were treated by oral gavage with UA (100 mg/kg) or vehicle every 2 days starting 7 days before IMQ challenge and throughout the experiment. SLE-like disease was induced by topical application of imiquimod to the postauricular skin (1.25 mg per ear) every 2 days for 6 weeks. At the endpoint, blood, urine, spleen, and kidneys were collected for ELISA, histology, immunofluorescence, and RT–qPCR analyses.

### ELISA

Plasma anti–double-stranded DNA (anti-dsDNA) antibody levels were measured by ELISA (MREDA, MF10351). Urinary albumin and creatinine were measured by ELISA (MREDA, M150631 and M132770, respectively). All assays were performed according to the manufacturers’ instructions. Urinary albumin-to-creatinine ratio (uACR) was calculated as urinary albumin divided by urinary creatinine.

### Histology, immunofluorescence, and clinical scoring

Psoriasis severity was assessed daily using a modified PASI scoring system (erythema, scaling, and thickness; each scored 0–4) as previously described ^52^. H&E staining and histopathological scoring were performed as previously described ^51^. For renal IgG deposition, kidney sections were stained with anti-IgG secondary antibody (Biodragon, BD9276) and counterstained with Hoechst 33342 (Beyotime, C1022). Fluorescence intensity was quantified by semiquantitative analysis as previously described ^53^, with identical acquisition settings applied across groups.

### Statistics

Statistical analyses were performed and graphs generated using GraphPad Prism 9. Data are presented as mean ± SD, with the exact format specified in each figure legend. For CCK-8 cell viability assays and luciferase reporter assays, differences among multiple treatment groups were assessed by one-way analysis of variance (ANOVA) followed by Dunnett’s post hoc test comparing each treatment group with the vehicle control. For qRT–PCR data and densitometric quantification of immunoblots, two-tailed unpaired Student’s t-tests were used. The number of biological replicates (n) for each experiment and the statistical tests applied are indicated in the corresponding figure legends. A *P* value < 0.05 was considered statistically significant.

### Study approval

All animal experiments were approved by the Institutional Animal Care and Use Committee of Peking University Health Science Center (approval number: DLASBE0119) and were performed in accordance with institutional guidelines.

## Supporting information

all supplemental data

## Data availability statement

The RNA-seq data generated in this study will be deposited in a public repository before journal publication. Other data supporting the findings of this study are available from the corresponding author upon reasonable request.

## Acknowledgments

We thank Dr. Tomas Lindahl (The Francis Crick Institute, United Kingdom), Dr. Yingyu Chen (Peking University, China), Dr. Zhengfan Jiang (Peking University, China), Dr. Fuping You (Peking University, China), Dr. Hong-Bing Shu (Wuhan University, China), for *Trex1^+/-^* mice, cell lines, and plasmids.

## Author contributions

SG conceived the study, designed and performed experiments, acquired and analyzed data, prepared figures, and wrote the manuscript. RCT provided input on experimental design and assisted with selected cell and molecular biology experiments. HY assisted with data analysis. AZ assisted with animal experiments. SSY assisted with plasmid and vector construction. YZ assisted with figure compilation and integration. LZ assisted with manuscript formatting. JZ conceived and supervised the study, provided overall guidance and resources, secured funding, and edited the manuscript. All authors reviewed and approved the final manuscript.

