## Supplementary material for "The gut microbiota metabolite Urolithin A mitigates JAK signaling to suppress cytokine–mediated autoimmune diseases": all supplemental data

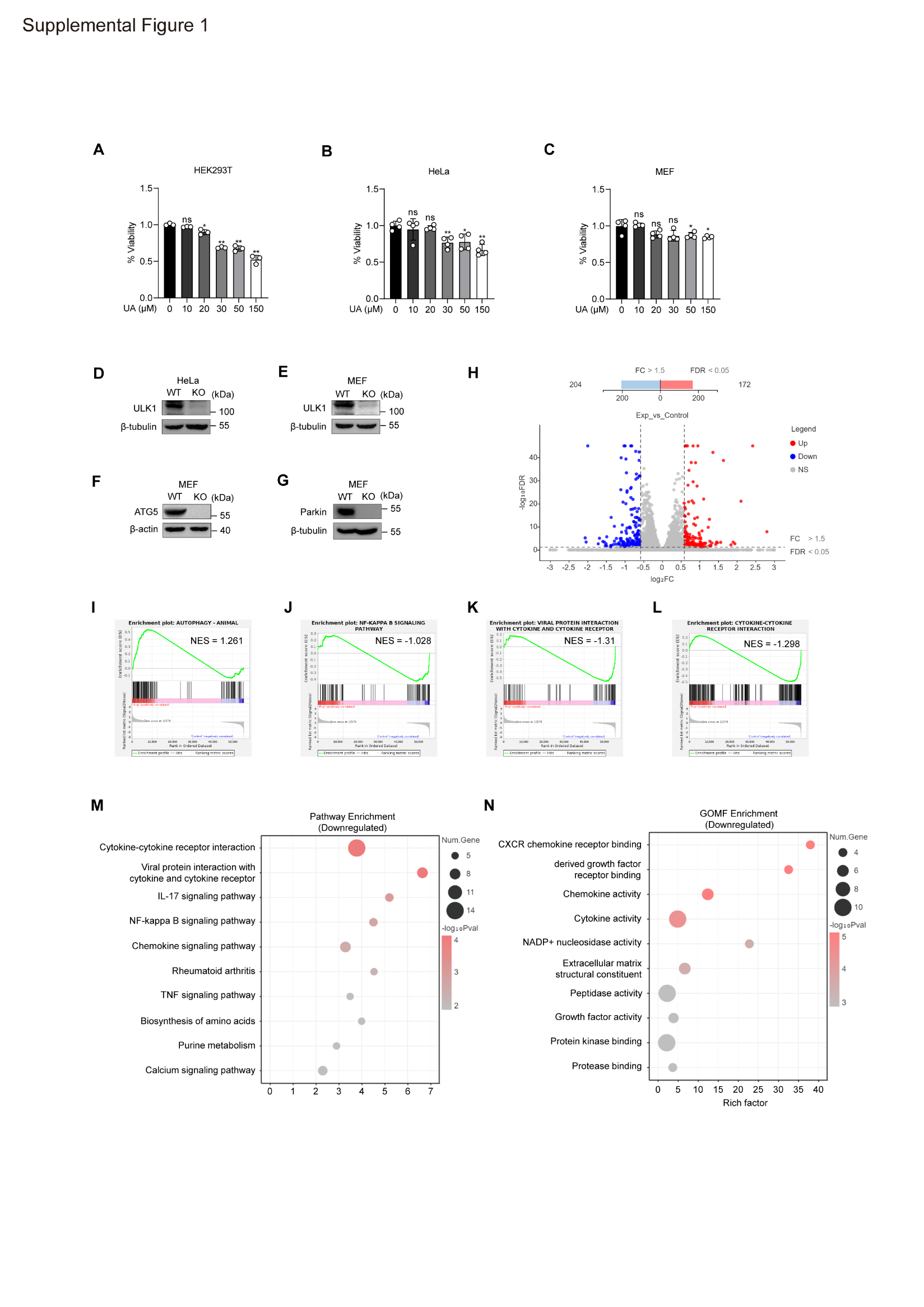

**Supplementary Figure 1 Cell viability of multiple cell lines, validation of knockout cell lines and extended transcriptomic analyses for UA-treated macrophages.**

**A**,**B**,**C** Cell viability of HEK293T, HeLa and MEF cells after 24 h exposure to UA at the concentrations indicated in the panel, measured by CCK-8 assay.

**D**,**E**,**F**,**G**, Immunoblot validation of genetic backgrounds used in Fig. 3: loss of *ULK1* in HeLa cells, *Ulk1* in MEFs, *ATG5* in MEFs and *Prkn* in MEFs.

**H**, Differential expression overview (volcano/MA plot) comparing UA-treated versus DMSO treated macrophages.

**I**,**J**.**K**.**L,** GSEA of UA-versus-vehicle-treated macrophages.

**M**,**N**, Integrated pathway and function readouts for downregulated genes.

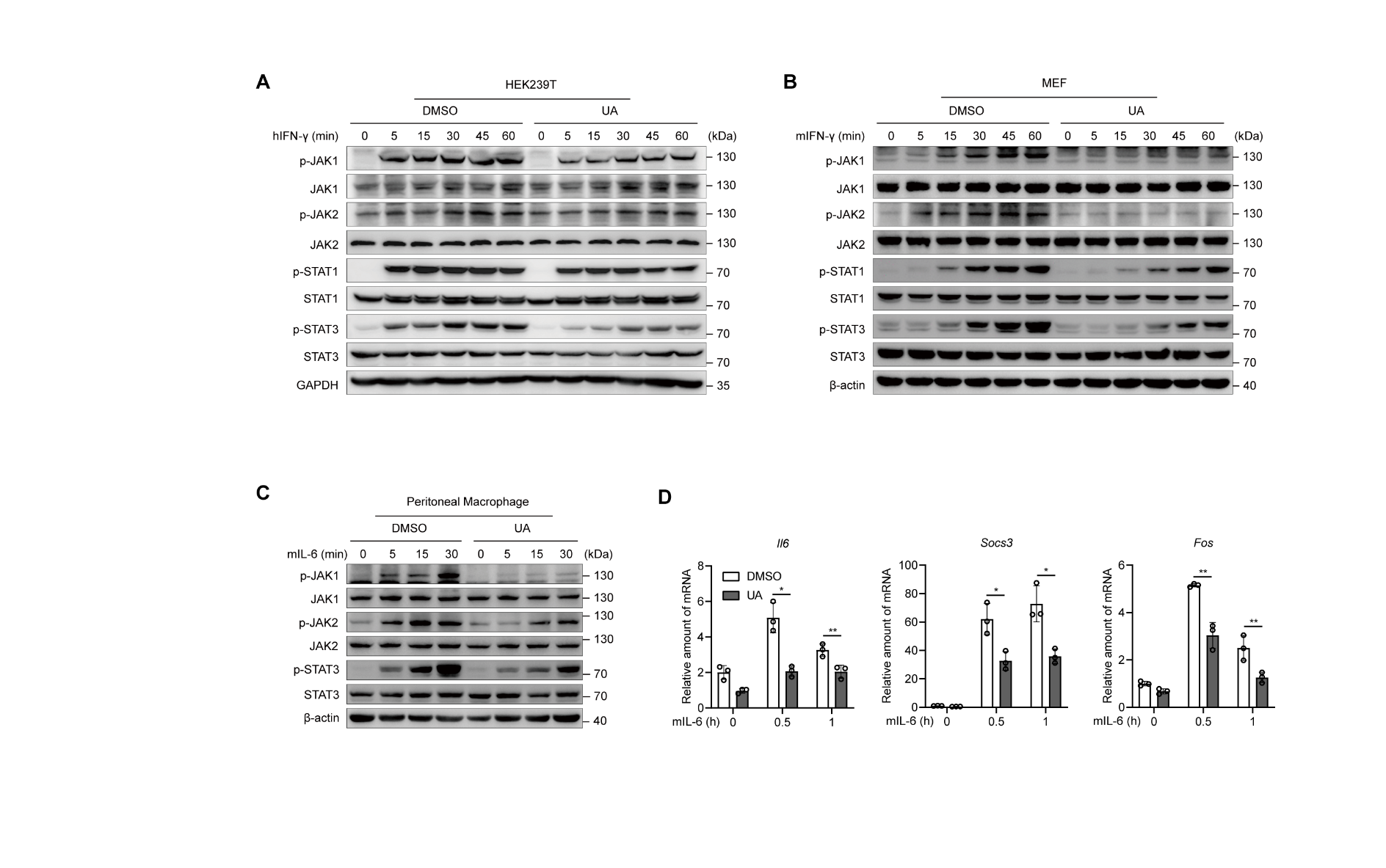

**Supplementary Figure 2 Supporting analyses for UA effects on IFN-γ and IL-6 signaling.**

**A**,**B**, Immunoblot confirmation of IFN-γ pathway inhibition by UA in HEK293T and MEF cells. Cells were pretreated with 10 µM UA for 24 h and stimulated with 10ng/mL IFN-γ as indicated.

**C**, Immunoblot confirmation of IL-6 pathway inhibition by UA in primary peritoneal macrophages pretreated with 10 µM UA for 24 h and stimulated with 10ng/mL mouse IL-6 as indicated.

**D**, qPCR analysis of *Il6*, *Socs3* and *Fos* mRNA levels in primary peritoneal macrophages pretreated with 10µM UA for 24 h and stimulated with 10ng/mL mouse IL-6 as indicated.

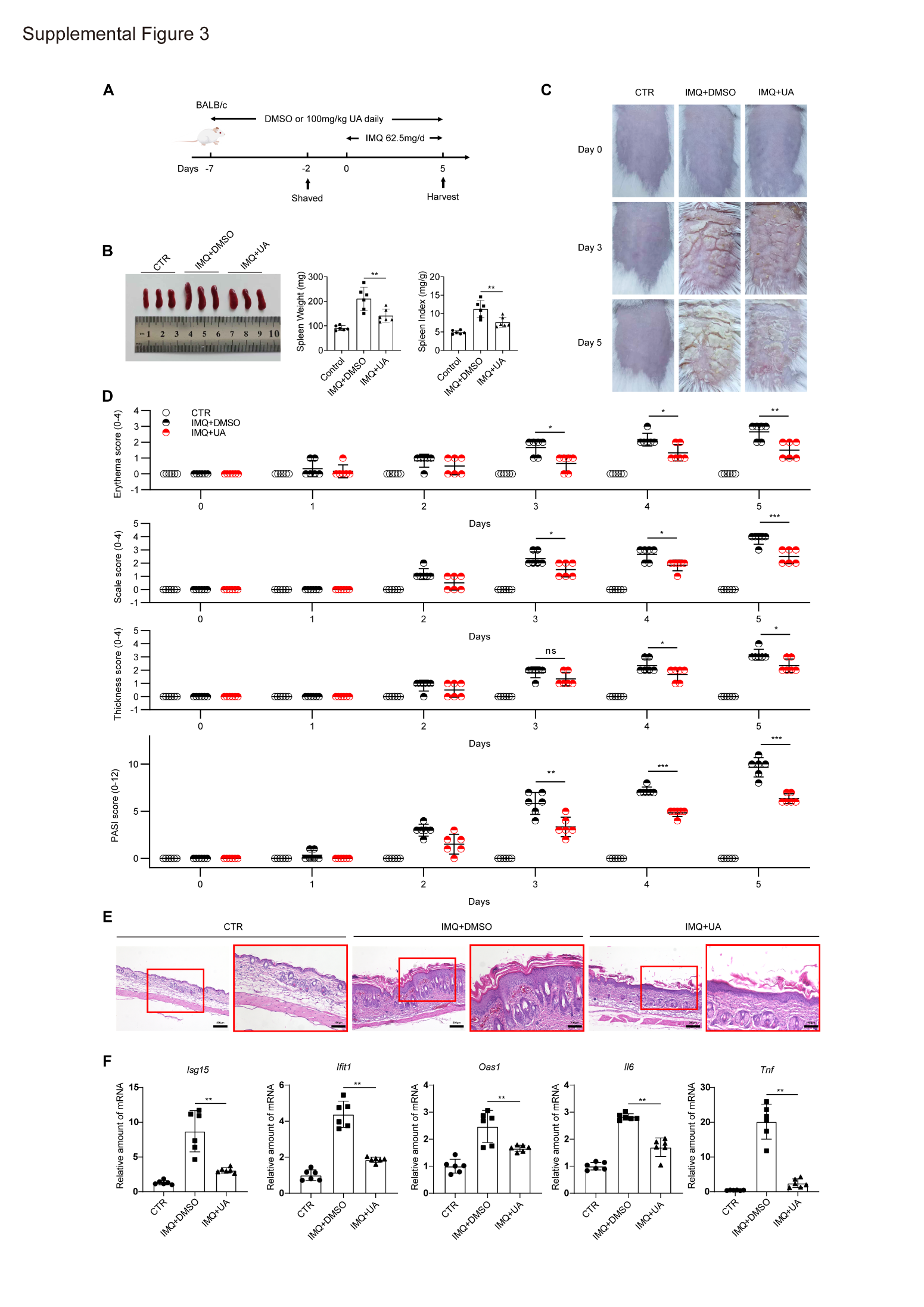

**Supplementary Figure 3 UA reduces cytokine and interferon-driven inflammation in an imiquimod-induced psoriasis-like model.**

**A**, Experimental scheme for topical imiquimod (IMQ)-induced psoriasis and concurrent oral UA or DMSO administration. (n = 6 mice per group)

**B**, Representative spleens, spleen weight and spleen index.

**C**,**D**, Representative skin appearance and quantification of erythema, scaling and thickness and PASI scores.

**E**, Histological analysis of dorsal skin showing decreased epidermal hyperplasia and inflammatory cell infiltration after UA treatment.

**F**, qPCR of lesional skin demonstrating reduced expression of interferon-stimulated and pro-inflammatory genes (*Isg15*, *Ifit1*, *Oas1*, *Tnf*, *Il6*) in UA-treated mice.

Statistical significance is indicated: ns = no significance, **P* < 0.05, ***P* < 0.01, ****P* < 0.001.

Supplemental Table 1. Antibodies for immunoblotting

| **Antibodies** | **Source** | **Identifier** | **Dilution** |
| --- | --- | --- | --- |
| Mouse anti-Flag | MBL | cat#M185-3L | 1:10000 |
| Mouse anti-HA | Abmart | cat#M20003S | 1:5000 |
| Mouse anti-GAPDH | Abmart | cat#M20006S | 1:5000 |
| Mouse anti-β-tubulin | Bioworld | cat#AP0064 | 1:5000 |
| Mouse anti-β-actin | Proteintech | cat#M30109S | 1:10000 |
| Rabbit anti-p-JAK1 (Tyr1034/1035) | Cell Signaling Technology | cat#3331S | 1:1000 |
| Rabbit anti-JAK1 | Selleck | cat#F0387 | 1:1000 |
| Rabbit anti-p-TYK2 (Tyr1054/1055) | Cell Signaling Technology | cat#9321S | 1:1000 |
| Mouse anti-TYK2 | Santa Cruz | cat#sc-5271 | 1:500 |
| Rabbit anti-p-JAK2 (Tyr1007/1008) | Abclonal | cat#AP0531 | 1:1000 |
| Rabbit anti-JAK2 | Abmart | cat#T55287 | 1:1000 |
| Rabbit anti-p-STAT1 (Tyr701) | Cell Signaling Technology | cat#9167S | 1:1000 |
| Rabbit anti-STAT1 | Cell Signaling Technology | cat#14995S | 1:1000 |
| Rabbit anti-p-STAT2 (Tyr690) | Cell Signaling Technology | cat#4441S | 1:1000 |
| Rabbit anti-STAT2 | Selleck | cat#F0713 | 1:1000 |
| Rabbit anti-p-STAT3 (Tyr705) | Cell Signaling Technology | cat#9145S | 1:1000 |
| Mouse anti-STAT3 | Cell Signaling Technology | cat#9193S | 1:1000 |
| Mouse anti-Parkin | Selleck | cat#F0296 | 1:1000 |
| Rabbit anti-ULK1 | Abmart | cat#T56902 | 1:5000 |
| Rabbit anti-ATG5 | Cell Signaling Technology | cat#12994T | 1:1000 |
| Mouse IgG | Sigma-Aldrich | cat#I5381 | 1:250 |
| Rabbit IgG | Sigma-Aldrich | cat#I5006 | 1:250 |
| HRP-conjugated anti-mouse IgG | Biodragon | cat#BF03001 | 1:8000 |
| HRP-conjugated anti-rabbit IgG | Biodragon | cat#BF03008 | 1:8000 |

Supplemental Table 2. Primers for qPCR

| **Primer name** | **Sequences** | |
| --- | --- | --- |
| Human *GAPDH* | Forward | ACCCACTCCTCCACCTTTGA |
|  | Reverse | CTGTTGCTGTAGCCAAATTCGT |
| Human *ISG15* | Forward | CTCTGAGCATCCTGGTGAGGAA |
|  | Reverse | AAGGTCAGCCAGAACAGGTCGT |
| Human *IFIT1* | Forward | GCCATTTTCTTTGCTTCCCCTA |
|  | Reverse | TGCCCTTTTGTAGCCTCCTTG |
| Human *OAS1* | Forward | CATCCGCCTAGTCAAGCACTG |
|  | Reverse | CCACCACCCAAGTTTCCTGTAG |
| Human *IL-6* | Forward | AGACAGCCACTCACCTCTTCAG |
|  | Reverse | TTCTGCCAGTGCCTCTTTGCTG |
| Human *SOCS3* | Forward | CGCCACTTCTTCACGCTCAG |
|  | Reverse | TCGGAGGAGGGTTCAGTAGGT |
| Human *FOS* | Forward | GCCTCTCTTACTACCACTCACC |
|  | Reverse | AGATGGCAGTGACCGTGGGAAT |
| Human *CXCL9* | Forward | TCTTGCTGGTTCTGATTGG |
|  | Reverse | AAGGATTGTAGGTGGATAGTC |
| Human *CXCL10* | Forward | CTCTAAGTGGCATTCAAGGA |
|  | Reverse | GGATTCAGACATCTCTTCTCA |
| Mouse *β-actin* | Forward | AGAGGGAAATCGTGCGTGAC |
|  | Reverse | CAATAGTGATGACCTGGCCGT |
| Mouse *Ifit1* | Forward | GAACCCATTGGGGATGCACAACCT |
|  | Reverse | CTTGTCCAGGTAGATCTGGGCTTCT |
| Mouse *Ifit2* | Forward | CGGAAAGCAGAGGAAATCAA |
|  | Reverse | TGAAAGTTGCCATACCGAAG |
| Mouse *Isg15* | Forward | CATCCTGGTGAGGAACGAAAGG |
|  | Reverse | CTCAGCCAGAACTGGTCTTCGT |
| Mouse *Oas1* | Forward | GCCTGATCCCAGAATCTATGC |
|  | Reverse | GAGCAACTCTAGGGCGTACTG |
| Mouse *Cxcl10* | Forward | GCCGTCATTTTCTGCCTCA |
|  | Reverse | CGTCCTTGCGAGAGGGATC |
| Mouse *Cxcl9* | Forward | GGAGTTCGAGGAACCCTAGTG |
|  | Reverse | GGGATTTGTAGTGGATCGTGC |
| Mouse *Il6* | Forward | TACCACTTCACAAGTCGGAGGC |
|  | Reverse | CTGCAAGTGCATCATCGTTGTTC |
| Mouse *Socs3* | Forward | GGACCAAGAACCTACGCATCCA |
|  | Reverse | CACCAGCTTGAGTACACAGTCG |
| Mouse *Fos* | Forward | GGGAATGGTGAAGACCGTGTCA |
|  | Reverse | GCAGCCATCTTATTCCGTTCCC |
| Mouse *Tnf* | Forward | GGTCCCCAAAGGGATGAGAA |
|  | Reverse | TGAGGGTCTGGGCCATAGAA |
